# Sensory Input, Sex, and Function Shape Hypothalamic Cell Type Development

**DOI:** 10.1101/2024.01.23.576835

**Authors:** Harris S. Kaplan, Brandon L. Logeman, Kai Zhang, Celine Santiago, Noor Sohail, Serhiy Naumenko, Shannan J. Ho Sui, David D. Ginty, Bing Ren, Catherine Dulac

## Abstract

Mammalian behavior and physiology undergo dramatic changes in early life. Young animals rely on conspecifics to meet their homeostatic needs, until weaning and puberty initiate nutritional independence and sex-specific social interactions, respectively. How neuronal populations regulating homeostatic functions and social behaviors develop and mature during these transitions remains unclear. We used paired transcriptomic and chromatin accessibility profiling to examine the developmental trajectories of neuronal populations in the hypothalamic preoptic region, where cell types with key roles in physiological and behavioral control have been identified^1–6^. These data reveal a remarkable diversity of developmental trajectories shaped by the sex of the animal, and the location and behavioral or physiological function of the corresponding cell types. We identify key stages of preoptic development, including the perinatal emergence of sex differences, postnatal maturation and subsequent refinement of signaling networks, and nonlinear transcriptional changes accelerating at the time of weaning and puberty. We assessed preoptic development in various sensory mutants and find a major role for vomeronasal sensing in the timing of preoptic cell type maturation. These results provide novel insights into the development of neurons controlling homeostatic functions and social behaviors and lay ground for examining the dynamics of these functions in early life.

## Introduction

Animals’ interactions with their internal and external environments change considerably as they develop. The maturation of sensory organs alters the detection of sensory inputs, such as vision after eye opening, and homeostatic needs such as sleep, thermoregulation and appetite are met in new ways as young animals mature and reach independence. Social relationships are transformed over time, for example with parents upon weaning and through the emergence of sex-specific reproductive and defensive behaviors after puberty. How these changes affect, or are affected by, the maturation of neuronal circuits is mostly unknown. Our current understanding of how brain circuits develop and mature mainly comes from studies of sensory system development. In visual and other sensory pathways, the emergence of distinct cell types and their spatial organization, connectivity, and activity patterns involve an interplay of genetic and activity-dependent mechanisms^7–10^. By contrast, despite decades of fierce debate on the roles of genetic and environmental information in the emergence of species- and sex-specific functions^11–13^, developmental mechanisms underlying the corresponding brain circuits are poorly understood. Distinct physiological and social functions emerge at various stages of postnatal development, yet how behavioral and homeostatic transitions are reflected in the development of specific neuronal populations remains unclear. In addition, despite clear evidence for a key role of environmental information in the proper maturation of social and survival behaviors^14–16^, how, where and when critical periods may shape the corresponding neuronal populations is still unknown^17^.

The preoptic area (POA) of the hypothalamus is an essential brain hub underlying the control of homeostatic functions and social behaviors. POA cell types identified by specific molecular signatures and residing in anatomically defined POA sub-regions have been associated with various physiological and social functions in adults. For example, VLPO Tac1+^3^ and Gal+^5^ neurons are involved in sleep/wake control, MnPO^Agtr1a^ neurons in thirst^2^, and MPN^Gal^ neurons in parenting^1,18,19^. The close proximity of cell types involved in such diverse survival and social functions may be conducive to mutual interactions between regulators of various needs^20^, and may reflect shared circuit architecture^6^. In early postnatal life, the coordinated regulation of homeostatic and social functions may be especially critical, as survival needs such as warmth and nutrient intake are dependent on interactions with conspecifics, usually parents or siblings. How and when these different neuronal populations emerge and mature remains unexplored.

One might expect brain areas involved in essential survival functions, such as the POA, to develop early in life and largely independently from environmental influences. However, key developmental steps such as the maturation of hypothalamic connectivity^21,22^ and myelination^23,24^ take place surprisingly late compared to other brain areas, consistent with protracted changes in functions like sleep, thermoregulation, and social interactions. Recent work has also shown that some hypothalamic circuits may be shaped by external inputs in early life^16,25^ or in adulthood^26^. However, the mechanisms, timing, and sensitivity to environmental influences of POA development and maturation have remained poorly understood and have, so far, lacked the cell type resolution obtained in studies of homeostatic and social functions in the adult. The POA also shows substantial molecular and functional sex differences^27–30^, yet the degree to which POA cell type maturation unfolds differently in males versus females remains underexplored.

Here, we use paired single-nucleus gene expression and chromatin accessibility sequencing (snRNA-seq and snATAC-seq) to map the developmental trajectories of 147 neuronal cell types in the POA and surrounding brain regions in mice. POA neurons are born between embryonic day 12 (E12) and E16 from progenitors lining the third ventricle, and they migrate radially to their ultimate sub-regional localization, with little to no migration into the POA from adjacent regions^31^. Recent work has described developmental transcriptomic changes in other hypothalamic areas^32^ or early embryonic stages in which cell migration and cell determination occur^33–35^. To explore further developmental processes within the POA during which cell differentiation leads to neuronal maturation and adult function, we performed snRNA-seq and snATAC-seq at E16, adult (postnatal day 65, P65), and six intermediate ages. To identify cell types with known homeostatic or social behavior functions, we mapped our data to a previously published molecular, spatial, and functional atlas of adult POA cell types^1^. We uncover diverse maturational trajectories that differ by POA sub-region, behavioral function, and sex, delineate the emergence of signaling networks involved in social or homeostatic functions, and identify key transcription factor drivers of these changes. Finally, we profile the POA of various mutants impaired in specific sensory modalities and identify a major age-specific role for vomeronasal input in ensuring normal POA developmental timing. Our findings provide novel insights into POA development and lay the foundation for a mechanistic understanding of critical periods in instinctive behavior circuit development and function in early life.

### A transcriptional atlas of POA cell type development

To build a transcriptional atlas of POA development, we performed droplet-based paired snRNA-seq and snATAC-seq at eight ages from E16 to P65 (Fig 1a). Ages were chosen to capture key transitions associated with birth, weaning, and puberty. At each age, we dissected the anterior portion of the mouse hypothalamus containing the POA and surrounding regions, pooling tissue from three brains per sample, and performed nuclei isolation and library preparation for two biological replicates per sex per age. After quality control filtering, we obtained snRNA-seq profiles from 206,413 nuclei, with snATAC-seq profiles for 198,753 of those nuclei.

**Fig. 1.**
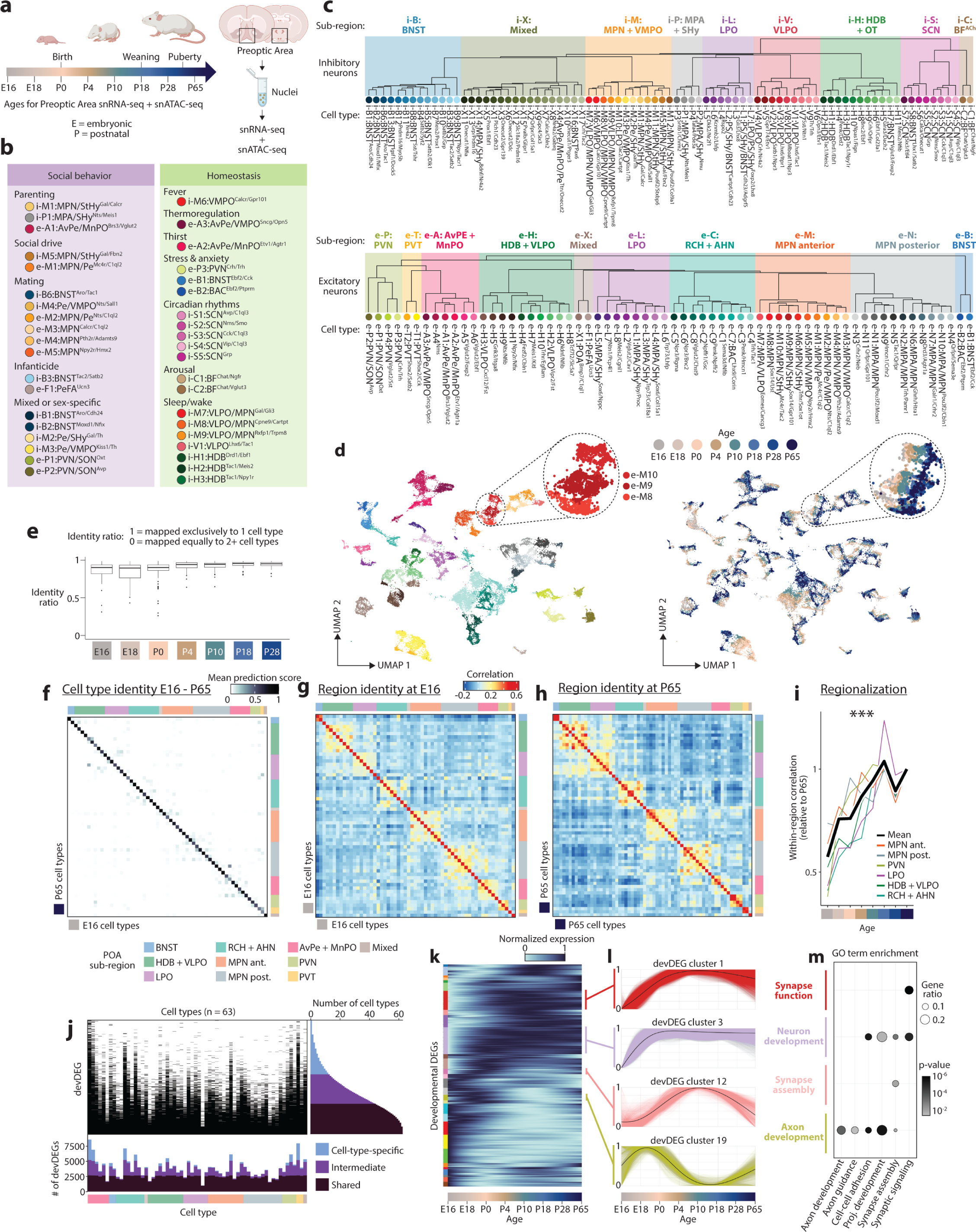
A molecular atlas of hypothalamic preoptic area cell types across development. **(a)** Schematic of experiments: preoptic area (POA) and surrounding regions were dissected, nuclei extracted, and snRNA-seq + snATAC-seq libraries prepared at 8 developmental stages, 2 male and 2 female samples per age. **(b)** Cell types with previously identified roles in social behavior or homeostatic control. **(c)** Hierarchical clustering of snRNA-seq clusters at P65, comprising 83 inhibitory (upper) and 64 excitatory clusters (lower), organized according to sub-regional identity (shading on tree). ACh, acetylcholinergic; AHN, anterior hypothalamic nucleus; AvPe, anteroventral periventricular nucleus; BF, basal forebrain; BNST, bed nucleus of the stria terminalis; HDB, horizontal limb of the diagonal band nucleus; LPO, lateral preoptic area; MnPO, median preoptic nucleus; MPA, medial preoptic area; MPN, medial preoptic nucleus; OT, olfactory tubercle; PVN, paraventricular hypothalamic nucleus; PVT, paraventricular thalamic nucleus; RCH, retrochiasmatic area; VLPO, ventrolateral preoptic nucleus; VMPO, ventromedial preoptic nucleus; SCN, suprachiasmatic nucleus. **(d)** UMAP across all 8 ages for excitatory clusters. Left, colored by cluster (key in **(c)**, lower); right, by age. Inset: gradient across age for 3 clusters. **(e)** Identity ratio quantifies how well clusters from young ages computationally map onto adult clusters: identity ratio of 1 denotes complete mapping to a single cluster, and identity ratio of 0 denotes equal mapping to 2 or more. Each data point represents a single cluster, averaged across all cells. **(f)** Mapping of E16 onto P65 clusters using canonical correlation analysis (excitatory neurons). Prediction score is averaged across cells within a cluster. **(g)** Correlation between pseudobulked gene expression of all clusters at E16. Colored bars on top and right indicate sub-region. **(h)** As in **(g)** but at P65. **(i)** Correlation between clusters within a sub-region, at each age, normalized to correlation at P65. ***p < 0.0001 for one-way ANOVA, effect of age.. **(j)** Differentially expressed genes (DEGs) were determined within each cluster across all ages (devDEGs). Right histogram: devDEGs are categorized according to the number of clusters in which they are identified: >70% (“shared”), <20% (“cell-type-specific”) or in between (“intermediate”). Bottom histogram: number of devDEGs for each cluster. **(k)** Normalized, cell-type-specific gene expression changes (fit with a trend curve) across all devDEGs, clustered (left colored bars) according to time course. **(l)** Four examples of devDEG clusters. Each colored line indicates the cell-type-specific gene expression change of a single devDEG; each black line indicates the mean across all devDEGs in the cluster. **(m)** GO term enrichment for the four examples of devDEG clusters shown in **(l)**.

To identify major cell classes among snRNA-seq profiles, we used a combination of Louvain clustering and gene marker analysis (Extended Data Fig 1a and Methods). After identifying neuronal and non-neuronal populations, we clustered neuronal profiles (150,180 nuclei) from adult (P65) samples, resulting in 83 GABAergic and 64 glutamatergic cell types. To validate our clustering approach and identify putative functional annotations for the corresponding cell types, we used label transfer^36^ to map the P65 cells onto our previously published scRNA-seq and MERFISH atlas of adult POA cell types^1^. This step revealed a high degree of correspondence (70% of cells mapping to the top matched cluster on average) between our de novo clusters and previously identified cell types, including those with known roles in homeostasis or social behavior (Fig 1b, Extended Data Fig 1b-c and Extended Data Table 1). In some cases (26 / 63 reference scRNA-seq clusters), individual clusters in the reference atlas were split into multiple clusters in our study, indicating increased resolution due to higher cell numbers and advances in sequencing technologies. Supervised hierarchical clustering based on correlation (Fig 1h) and spatial information from MERFISH (inferred from label transfer) revealed that cell types localized in a given POA sub-region share transcriptional signatures. We therefore defined a two-tier classification, with the first tier corresponding to sub-regional identity and the second tier corresponding to cell type (Fig 1c and Supplementary Table 1). Cell type IDs denote (1) sub-regional group (e.g. i-M1-12 for 12 inhibitory MPN or e-P1-4 for 4 excitatory PVN cell types), (2) specific sub-regional localization if known (e.g. i-M1:MPN/StHy), and (3) marker genes based on both the literature and this study (e.g. i-M1:MPN/StHy^Gal/Calcr^). Some cell types did not map to the published POA atlas, likely falling outside of the MERFISH imaging area; we assessed their localization by performing in situ hybridization with a panel of cell-type-specific genes (Extended Data Fig 2).

Next, we asked how snRNA-seq profiles from POA of younger animals map onto adult cell types. Using a similar label transfer approach^36^ (Methods), we sequentially mapped datasets from each developmental stage onto the cluster identities obtained from older stages (from P28 to P65, then P18 to P28+P65, etc.) (Fig 1d-e and Extended Data Fig 3 and 4a). This approach was able to identify nearly all adult clusters at each younger age, even at the earliest age E16 (Fig 1e-f and Extended Data Fig 4b), indicating that nearly the full complement of cell types is diversified well before birth. While nearly all clusters were identifiable at all ages, each cluster appeared in UMAP space as a temporal gradient of cells ordered across age (Fig 1d inset and Extended Data Fig 4a inset). This suggests the existence of gradual maturational changes in gene expression occurring on top of stable cell type identities.

We next examined paired snATAC-seq libraries, using cell type identities from snRNA-seq clustering. To identify candidate transcriptional drivers of neuronal identity, we determined cell-type-specific genomic loci of chromatin accessibility and transcription factor motifs enriched in those loci (Extended Data Fig 5a-b). Among this rich dataset, both known and novel candidate transcriptional regulators were identified. For example, POU-domain containing transcription factor motifs were identified as enriched in PVN neurosecretory cell types^37^. The motif for the transcription factor family ROR/REV-ERB was highly enriched in two SCN cell types, i-S1:SCN^Avp/C1ql3^ and i-S3:SCN^Cck/C1ql3^, which adds cell-type-specificity to a previously identified role for REV-ERB in SCN control of circadian insulin sensitivity in humans^38^. The estrogen receptor Esr1 was enriched in cell types with known sex differences in function or cell number^1,6^, also further examined in a later section (Fig 4). This dataset thus represents a precious atlas of candidate transcriptional regulators determining cell type identity.

We next asked how regional identity signatures change during development. In the adult, clusters belonging to the same POA sub-region were more likely to be correlated transcriptionally (Fig 1h and Extended Data Fig 4d). Correlation maps at E16 revealed within-region correlations resembling those at P65, albeit less defined (Fig 1g-h and Extended Data Fig 4c-d); with age, these correlations gradually strengthened, especially between P0 and P10 (Fig 1i and Extended Data Fig 4e). Each sub-region expressed a specific set of marker genes, many of which showed changes in region specificity with age (Extended Data Fig 4f). Sub-region-specific marker gene expression also came to gradually resemble adult patterns with age, especially between P0 and P10 (Extended Data Fig 4g). We confirmed these findings for specific genes and sub-regions using in situ hybridization (Extended Data Fig 2). Altogether, our data show that, between P0 and P10, the POA undergoes a regionalization process by which cell types within a given sub-region adopt common transcriptional signatures specific to that sub-region, similar to the dynamic arealization process occuring in cortex^39^. Our marker gene analysis also identified a set of regional transcription factors that largely maintain their expression and regional specificity from E16 to P65 (Extended Data Fig 4h). Several of these transcription factors also show a region-specific enrichment of their motifs in chromatin accessibility data, thus potentially orchestrating regional identity. These include Tcf7l2 motifs enriched in PVT and HDB/VLPO neurons, Eomes motifs enriched in BNST excitatory neurons, Sox6 motifs enriched in BNST inhibitory neurons, and Pou3f motifs enriched in PVN, BF, and MPA/SHy neurons (Extended Data Fig 5c).

We next examined developmental changes in gene expression at the cell type level. For each individual cell type, we identified a set of genes that are differentially expressed between ages (devDEGs). Some genes were identified as devDEGs across multiple clusters, indicative of shared neuronal maturational changes, while others were identified as devDEGs in only one or a few clusters (Fig 1j). Accordingly, we classified each devDEG as shared, cell-type-specific, or intermediate depending on the number of cell types in which it was identified. This revealed a set of ∼2500 devDEGs shared across nearly all cell types and up to 5000 additional devDEGs with various degrees of cell type specificity (Fig 1j). To examine developmental timing, we fit trends to devDEG changes across age using a generalized additive model, pooled devDEG trends across cell types, and clustered them according to their dynamics^40^ (Fig 1k). Some devDEG clusters represented genes that showed increasing or decreasing expression at specific ages, while others showed more complex bimodal patterns (Fig 1l). Gene ontology analysis revealed devDEG clusters enriched in processes relevant for neuronal development (Fig 1m), indicating age-specific changes in functions such as axon guidance, cell adhesion, synapse assembly and synaptic signaling.

In summary, our dataset establishes a multiomic atlas of POA development comprising 147 neuronal populations distributed among 20 sub-regions, reveals a dynamic regionalization process, identifies candidate transcription factors regulating region and cell type identity, and describes cell-type-specific gene expression changes pertinent to the maturation of neuronal circuits.

### POA cell types follow diverse maturational trajectories

Homeostatic controls and social behaviors change significantly and along various timelines postnatally. For example, mice begin to independently thermoregulate around P10 and eat around P18, whereas mating and aggressive behaviors are typically not displayed until after P28 (puberty). We therefore asked whether distinct POA cell types display specific maturation trajectories. For each cell type, we quantified its distance^41^ between each pre-adult age and adult (Fig 2a). This analysis revealed a wide diversity of maturational trajectories. For example, i-C1:BF^Chat/Ngfr^, a cholinergic basal forebrain cell type involved in arousal, is by far the most mature cell type at P0-P10, essentially already adult-like by P10 (Fig 2b) (see Extended Data Table 1 for all subsequent citations for social and homoeostatic functions). In contrast, e-P3:PVN^Crh/Trh^, which initiates the corticosterone stress hormone response, is still relatively immature until after P28. Further, e-P3:PVN^Crh/Trh^ exhibits a stepwise trajectory, maturing at two key stages: P10-P18 and P28-P65. Remarkably, these two stages showed the most maturation across cell types (Extended Data Fig 6a), which suggests that weaning (∼P16-P21) and puberty (∼P28-P40) are key developmental events in the POA. Another late-maturing cell type is i-M1:MPN/StHy^Gal/Calcr^, which has well-studied roles in parenting behavior. However, these two late-maturing cell types showed very different maturational timing: in contrast to the non-linear, stepwise maturation of e-P3:PVN^Crh/Trh^, i-M1:MPN/StHy^Gal/Calcr^ exhibits a gradual maturation pattern, with maturation occurring at every age (Fig 2b).

**Fig 2.**
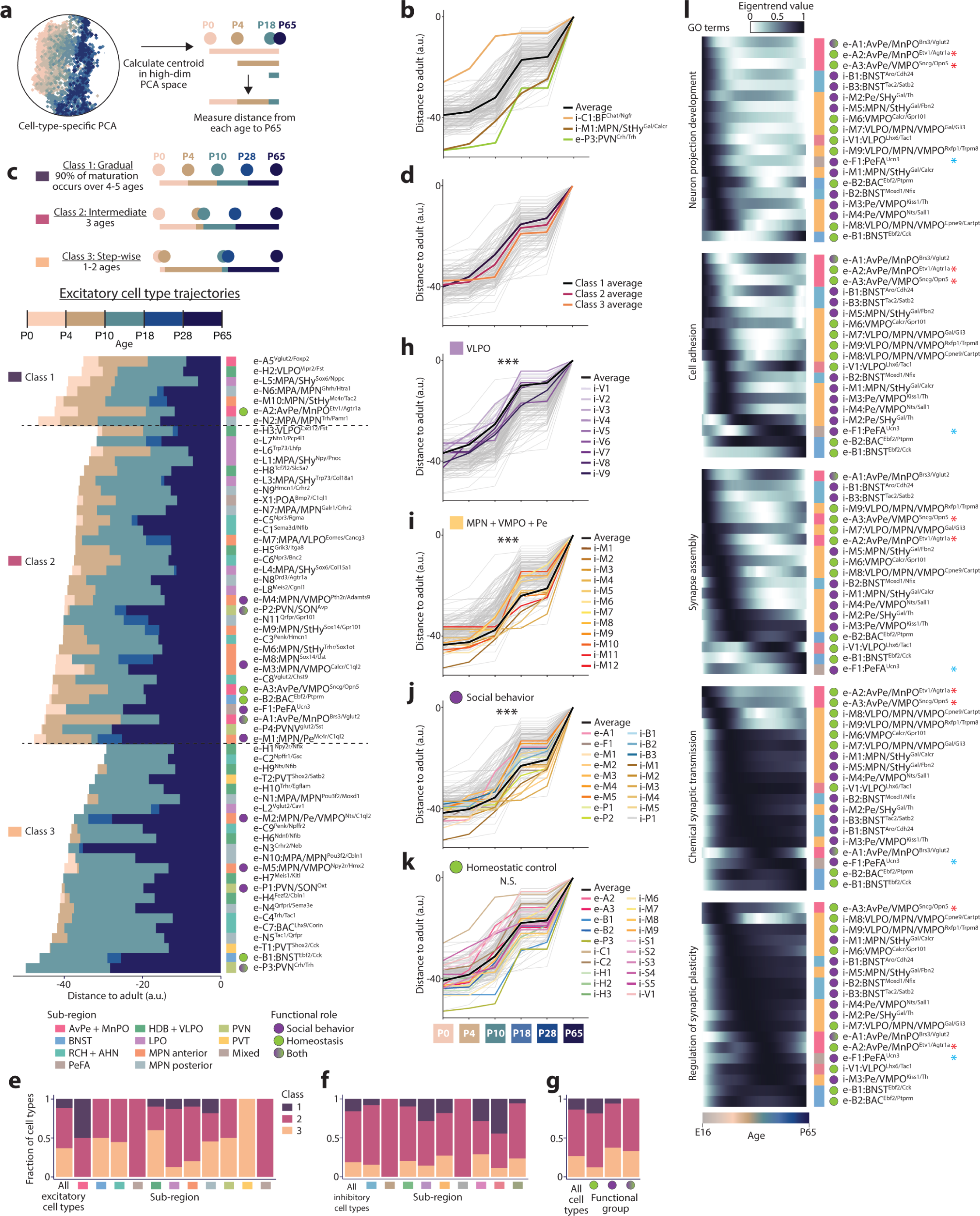
Developmental trajectories of POA cell types. **(a)** Strategy for quantifying maturity as distance between each age’s centroid and the adult centroid in PCA space, calculated separately for each cell type. **(b)** Distance trajectories across age for all cell types (gray), average (black) and extreme cell types of interest (colored). **(c)** Upper: schematic displaying different trajectory Classes. Lower: Distance trajectories across excitatory cell types. Trajectories were split into three classes depending on whether 90% of the maturation occurs across 4-5 ages (Class 1: Gradual), 3 ages (Class 2: Intermediate), or 1-2 ages (Class 1: Stepwise). **(d)** Distance trajectories averaged across each class. **(e)** Fractional class distribution of excitatory cell types, split by sub-region. **(f)** As in **(e)** but for inhibitory cell types. **(g)** Fractional class distribution of all cell types or split by functional group. **(h-i)** Distance trajectories for inhibitory VLPO **(h)** or MPN + VMPO + Pe **(i)** cell types. ***p < 0.0001 for two-way ANOVA, effect of cell type subset. **(j-k)** Distance trajectories for cell types with roles in social behavior **(j)** or homeostatic control **(k)**. ***p < 0.0001 or Not Significant (N.S.) for two-way ANOVA, effect of cell type subset. **(l)** Eigentrend values summarize expression for devDEGs in select GO terms across age, for clusters of interest. Red and blue asterisks indicate clusters referred to in text as tending to show early (red) or late (blue) changes.

To better monitor these distinct trajectories, we examined them along two dimensions: the mode of maturation, i.e gradual vs. stepwise (Fig 2c-g and Extended Data Fig 6b-e) and the timing of maturation, i.e early vs. late (Fig 2h-k). We then asked whether the maturational trajectories along these dimensions correlated with features such as neurotransmitter, POA sub-region, or behavioral function. To distinguish gradual vs. stepwise maturation, we classified each cell type’s trajectory according to whether 90% of the maturational change occurred within 4-5 developmental stages out of 5 possible stages (gradual, Class 1); 3 stages (intermediate, class 2); or just 1-2 stages (stepwise, class 3) (Fig 2c for excitatory cell types and Extended Data Fig 6b for inhibitory cell types). Strikingly, nearly all cell types that matured in a stepwise manner (class 3) displayed dominant P10-P18 and P28-P65 steps (Fig 2c-d and Extended Data Fig 6b). Glutamatergic cell types tended to mature in a more stepwise fashion than GABAergic cell types (Fig 2c,e-f and Extended Data Fig 6b). Maturation class also varied by regional identity or functional role; for example, AvPe/MnPO cell types tended to mature gradually (despite being excitatory) whereas cell types involved in social behavior tended to have more stepwise trajectories (Fig 2e-g). Some POA sub-regions showed strong biases toward early or late maturation. Cell types in VLPO and AvPe/MnPO, two sub-regions involved in homeostatic functions like sleep, thirst, and thermoregulation, were typically more mature than other cell types, at all ages (Fig 2h and Extended Data Fig 6c). In contrast, cell types in MPN/VMPO/Pe and PVN, sub-regions involved in social behavior, were typically less mature than most other cell types, at all ages until adulthood (Fig 2i and Extended Data Fig 6d). Across all sub-regions, cell types involved in social behavior tended to be late maturing (Fig 2j), while cell types involved in homeostatic control showed a wide range of maturation timing (Fig 2k). Trajectory dependence on sub-region and function was corroborated by an alternative measure of maturation, in which the proportion of adult nearest neighbors was quantified at each younger age^40^ (Extended Data Fig 6e).

To assess differences in cell type maturation for genes associated with specific neurodevelopmental functions, we focused on devDEGs with ontology terms of interest. This revealed expected dynamics across most cell types, such as a progression from neuron projection development to cell adhesion and synapse assembly to synaptic function, as well as cell-type-specific deviations from the norm (Fig 2l). For example, consistent with the early maturation of AvPe/MnPO cell types (Extended Data Fig 6c), the e-A2:AvPe/MnPO^Etv1/Agtr1a^ cell type involved in thirst and the e-A3:AvPe/MnPO^Sncg/Opn5^ cell type involved in thermoregulation showed the earliest peaks in several gene categories (Fig 2l, red asterisks). In contrast, e-F1:PeFA^Ucn3^, which plays a key role in infanticidal behavior, and e-B1:BNST^Ebf2/Cck^, involved in anxiety and feeding behavior, maintain expression of cell adhesion and synapse assembly genes unusually late, suggesting synaptic changes well beyond most other cell types (Fig 2l, blue asterisks). Overall, our data reveal a great diversity in cell-type-specific maturational trajectories, which correlate in part with the cell type sub-regional identity and functional role.

### Developmental changes in POA signaling networks

In adults, POA neurons affect behavior through intra-hypothalamic and brain-wide signaling networks^3,4,6,19,42^. To examine when and how these signaling networks emerge and change during development, we quantified developmental gene expression changes in neuronal signaling systems, including neurotransmitters, monoamines, neuropeptides, and hormones.

GABA and glutamate synthesis and vesicle packaging enzymes were already highly expressed at E16 and showed little change with age (Extended Data Fig 7a). Several GABA and glutamate receptor genes, as well as genes for synaptic release machinery were expressed at E16 (Extended Data Fig 7b-c), suggesting that neurotransmitter signaling in POA could occur as early as E16. The expression of both GABA and glutamate receptor genes steeply increased from E16 to P4; after P4, overall levels (aggregated across genes and cell types) remained constant through adulthood (Fig 3a). Some individual receptor genes, such as the NMDA receptor subunit Grin2a, important for plasticity and neuronal maturation^43^, resembled changes to overall receptor levels (Fig 3b), whereas others, such as the AMPA receptor Gria3, showed later and more cell-type-specific dynamics (Fig 3c). These results suggest two distinct phases of neurotransmitter receptor expression dynamics: (1) a global increase between E16 and P4, which we interpret as an Establishment Phase, and (2) a reorganization of cell-type- and receptor-specific gene expression patterns between P4 and P65, viewed as a Refinement Phase.

**Fig 3.**
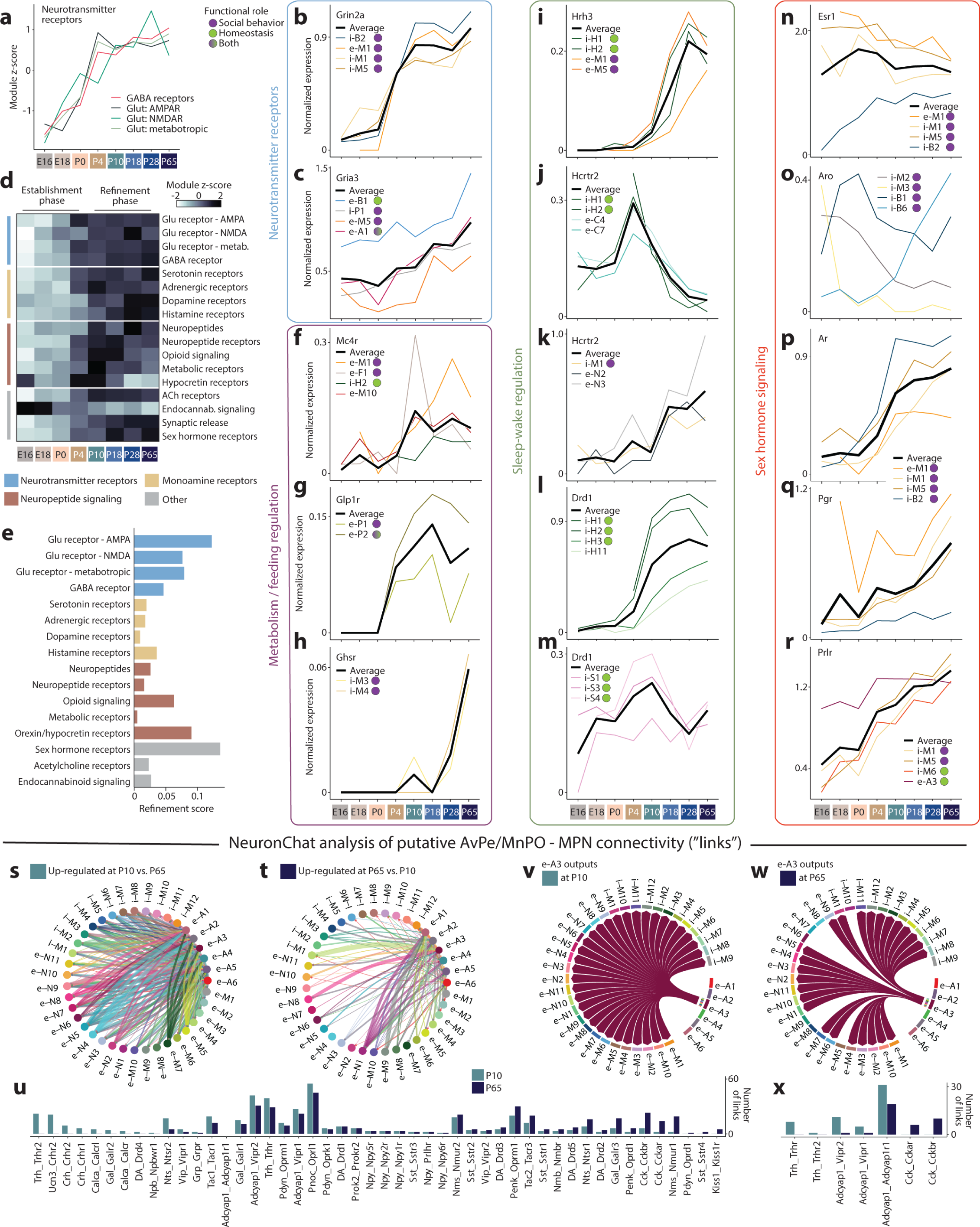
Maturational changes in POA neuronal signaling. **(a)** Module score aggregating expression across neurotransmitter receptor gene categories, then z-scored across age. **(b-c)** Log-normalized gene expression for glutamate receptor subunits Grin2a **(b)** or Gria3 **(c)**, averaged within each cluster across age, for example clusters (colored lines). Black line shows average across example clusters. **(d)** Module score aggregating expression across various signaling gene categories, then z-scored across age. **(e)** Refinement score quantifies the number of genes in each category that show significant changes between P10 and P65 (Refinement Phase), normalized by the number of genes in each category. **(f-h)** As in **(b-c)** for receptors involved in metabolism and feeding: Mc4r **(f)**, Glp1r **(g)**, and ghrelin receptor Ghsr **(h)**. **(i-m)** As in **(b-c)** for receptors involved in sleep/wake regulation: Hrh3 **(i)**, Hcrtr2 with example cell types involved in sleep/wake **(j)** or social behavior **(k)**, and Drd1 with example cell types in BF **(l)** or SCN **(m)**. **(n-r)** As in **(b-c)** for genes involved in sex hormone signaling: Esr1 **(n)**, aromatase **(o)**, Ar **(p)**, Pgr **(q)**, and Prlr **(r)**. **(s-t)** NeuronChat analysis of putative neuropeptide and monoamine connectivity between cell types in AvPe/MnPO and MPN at P10 versus P65. More links up-regulated at P10 **(s)** than at P65 **(t)**. **(u)** Number of NeuronChat neuropeptide and monoamine links identified at P10 or P65, sorted from left to right according to up-regulation with age. **(v-w)** Output links from e-A3 at P10 **(v)** or P65 **(w)**. **(x)** As in **(u)** but for e-A3 outgoing links only.

Next, we examined changes in other signaling systems, including monoamines, neuropeptides, and hormones. Nearly all gene sets showed overall patterns similar to neurotransmitter receptor genes, with an Establishment Phase during which overall expression levels increased from E16 to P4, followed by a plateau from P10 to P65 (Fig 3d). Notable exceptions to these trends include neuropeptides and dopamine receptors, which increased expression mostly between P4 and P10, and histamine receptors, which increased expression gradually until P18. Endocannabinoid signaling followed a different pattern altogether (Fig 3d): the cannabinoid receptor Cnr1 was already expressed at high, adult-like levels in most cell types at E16 (Extended Data Fig 7d), suggesting that endocannabinoid signaling may be particularly important in developing POA networks as it is in other brain regions^44,45^. Between P10 and P65, most signaling systems showed constant overall expression, yet individual genes continued to change expression in particular cell types, consistent with a Refinement Phase across many signaling systems (Extended Data Fig 7e-r). To better assess these changes, we quantified a Refinement Score for each signaling system, a measure of how many devDEGs were found between P10 and P65 (Extended Data Fig 7f). This analysis revealed that monoamine and neuropeptide signaling showed relatively low levels of refinement, compared to neurotransmitter receptors, opioid signaling, orexin/hypocretin signaling, and sex hormone receptors (Fig 3e), consistent with protracted changes in functions such as sleep or social behavior.

We next examined signaling systems with well-established roles in metabolism and appetite regulation, sleep/wake control, and social behavior, all of which show major postnatal changes. The POA receives significant input from Agrp+ and Pomc+ neurons in the Arcuate nucleus^46^. Arcuate neurons are regulated by circulating leptin and ghrelin through their cognate receptors Lepr and Ghsr and they exert their function in controlling food intake partly through release of neuropeptides that bind to the receptor Mc4r^21^. Mc4r expression is detectable in seven POA cell types, including e-M1:MPN/Pe^Mc4r/C1ql2^, which controls social drive in female mice, and e-F1:PeFA^Ucn3^, which regulate infanticide. In all seven cell types, Mc4r expression increased at P10 (Fig 3f and Extended Data Fig 7g). The leptin receptor Lepr is also expressed in e-M1:MPN/Pe^Mc4r/C1ql2^, as well as in e-A1:AvPe/MnPO^Brs3/Vglut2^, which has reported roles in parenting behavior and body temperature regulation. Lepr showed expression increases in these and other cell types with a similar timing as Mc4r (Extended Data Fig 7h). This suggests that peripheral (leptin) and Arcuate regulation of POA cell types, including cell types involved in social behavior, may begin between P4 and P10. This timing coincides with both the maturation of Arcuate projections^21^ and a surge in circulating leptin, shown to be critical for the establishment of Arcuate neuronal circuitry^47^. Prior to the development of Arcuate projections, the brainstem is a prominent regulator of feeding behavior, through projections to PVN already present at birth^21^. In adults, Glp1 signaling from brainstem to Glp1r+ neurons in PVN regulates food intake^48^. Consistent with an earlier regulation of feeding by brainstem compared to Arcuate nucleus, we find Glp1r expression in PVN is already high at P4 (Fig 3g). Finally, expression of the ghrelin receptor Ghsr was enriched in two POA cell types involved in mating behavior, including Kiss1+ neurons (Fig 3h). Ghsr expression increased particularly late in these two cell types, between P18 and P65. Altogether, these data indicate three developmental phases of metabolic control on POA cell types, marked by the onset of distinct receptor expression patterns: (1) brainstem regulation of PVN via Glp1 at P4; (2) Arcuate and peripheral (leptin) regulation of various POA cell types, including cell types involved in thermoregulation and social behavior, at P10; (3) ghrelin regulation of periventricular cell types involved in social behavior at P28.

Young animals show major changes in sleep and circadian rhythms, from early postnatal life and extending well into adolescence. We therefore examined developmental changes in orexin/hypocretin and histamine signaling. The histamine receptor Hrh3 showed no detectable expression until P4, followed by large increases from P10 to P28 (Fig 3i). These changes occurred in neurons in the HDB, part of the basal forebrain, a key brain region in arousal control. Unexpectedly, Hrh3 was also enriched in MPN neurons with roles in social drive and mating and showed similar changes with age, suggesting dynamic regulation of social behavior by sleep/wake signaling systems (Fig 3i). Such regulation was also apparent in hypocretin receptor expression: Hcrtr1 and Hcrtr2 were expressed in MPN neurons with well-established roles in parenting and social drive (Fig 3k), as well as basal forebrain neurons with known roles in sleep and arousal, including cholinergic neurons (Fig 3j). However, unlike Hrh3, Hcrtr1 and Hcrtr2 expression showed distinct developmental dynamics in cell types involved in social behavior (increased expression at P18-P28 (Fig 3k)) compared to cell types involved in sleep/wake control(a surprisingly transient peak of expression at P4-P10 (Fig 3j and Extended Data Fig 7i-j)). Hypocretin signaling may therefore play distinct roles, via different sets of cell types, at P4-P10 versus later ages. Like metabolic signals, sleep/wake signals are poised to affect social behavior, via histamine and hypocretin signaling in MPN cell types particularly later in life.

Dopamine signaling in the hypothalamus is best studied for its role in social behavior^28,49^. Dopaminergic neurons in various brain regions are also involved in sleep/wake control^50^. Consistent with both roles, we found dopamine receptor expression enriched in sub-regions involved in social behavior (BNST, MPN) and sleep/wake control (basal forebrain, SCN) (Extended Data Fig 7k). Dopamine receptors exhibited complex receptor- and cell-type-specific developmental dynamics in BNST and MPN cell types (Extended Data Fig 7l), including expression as early as E16. We identified two related cell types that express Th, Ddc, and Slc18a2 and are thus poised to synthesize, package, and release dopamine, and which play key roles in mating and parenting behavior^1,28,49^; expression of these genes were present at low levels at E16 and increased with age (Extended Data Fig 7m). Dopamine receptor expression in basal forebrain and SCN showed region-specific differences in developmental dynamics. Drd1 and Drd3 were both expressed in basal forebrain cell types involved in sleep/wake regulation, but only from P10 onwards (Fig 3l and Extended Data Fig 7l). In contrast, Drd1 expression in SCN cell types that dictate circadian rhythms was present as early as E18 and constant through to adulthood (Fig 3m). In summary, dopamine receptor expression shows highly region- and cell-type-specific developmental dynamics, indicating complex regulation of monoamine signaling across age.

Next, we focused on sex hormone signaling, which drives sexual differentiation in the POA both at puberty (P28 to P65) and perinatally (E18-P4), when a transient male-specific surge in testosterone impacts gene expression following conversion to estrogen by the enzyme aromatase^29,51^. Consistent with this model, the estrogen receptor Esr1 was enriched perinatally in MPN/Pe and BNST, two regions with well-established sex differences in gene expression^27,29^ (representative example populations shown in Fig 3n-r; others shown in Extended Data Fig 7o-r). Esr1 expression was largely maintained in those cell types through adulthood and showed small increases in additional cell types (Fig 3n and Extended Data Fig 7o). Estrogen signaling in the male brain has been proposed to occur in a paracrine manner: testosterone is converted to estrogen in a small number of aromatase-expressing cell types, followed by estrogen binding to Esr1 in a larger number of Esr1-expressing cell types^1^. Indeed, we observed aromatase expression at birth in three cell types: i-B1:BNST^Aro/Cdh24^, i-M2:Pe/SHy^Gal/Th^, and i-M3:Pe/VMPO^Kiss1/Th^ (Fig 3o). Intriguingly, all three showed a decrease in aromatase expression with age, whereas a fourth cell type, i-B6:BNST^Aro/Tac1^, showed an age-dependent increase in aromatase expression (Fig 3o and Extended Data Fig 7n). This suggests that distinct sets of neurons convert testosterone to estrogen at birth compared to adulthood. Further, i-M2:Pe/SHy^Gal/Th^ and i-B6:BNST^Aro/Tac1^ did not express detectable levels of Esr1, consistent with paracrine estrogen signaling. In contrast to Esr1, expression of the testosterone receptor Ar, the progesterone receptor Pgr, and the prolactin receptor Prlr is low perinatally and showed substantial increases with age, in a larger number of cell types (Fig 3p-r and Extended Data Fig 7p-r). Consistent with a role in physiological changes during pregnancy and nursing^52^, Prlr was expressed in cell types involved in homeostatic control functions, including fever, sleep, and thermoregulation (Fig 3r and Extended Data Fig 7r).

In summary, our analysis reveals complex expression dynamics in the development of POA signaling, such that distinct signaling networks emerge and may act at different ages. Moreover, our data suggests substantial crosstalk between signaling systems and cell types involved in homeostasis and social behavior.

POA neurons are highly peptidergic, with individual cell types typically expressing a dozen or more distinct neuropeptides. To understand the maturation of peptide signaling networks, we used NeuronChat^53^, a software package that infers neuronal connectivity based on statistical testing of pairs of peptide and receptor expression (e.g. Vip-Vipr1), termed links. We focused this analysis on connectivity between MPN, a region largely but not exclusively involved in social behavior, and AvPe/MnPO, which includes cell types associated with key homeostatic functions like thermoregulation and thirst. These two regions are known to project to one another^4,19,54^, with substantial connectivity already documented at P10^22,55^. Strikingly, NeuronChat identified denser putative inter-connectivity at P10 than at P65 (Fig 3s-t) with multiple neuropeptide links only found at P10 but not P65, such as Crh-Crhr1/2, Gal-Galr2, and Calca-Calcr (Fig 3u). Few links were found at P65 but not at P10, such as Kiss1-Kiss1r associated with reproduction. These changes occurred across several cell types (Extended Data Fig 7s-x); for example, e-A3:AvPe/MnPO^Sncg/Opn5^, an important player in thermoregulation, showed output links to all MPN cell types at P10, largely due to ubiquitous expression of the Adcyap1r1 receptor in MPN (Fig 3v-x). At P65, however, Adcyap1 signaling from e-A3:AvPe/MnPO^Sncg/Opn5^ was reduced and Trh signaling was lost; as a result, several MPN cell types no longer received putative input from e-A3:AvPe/MnPO^Sncg/Opn5^ (Fig 3v-x). These changes may indicate both pruning of exuberant connectivity between P10 and P65, as well as extensive neuropeptidergic signaling in developmental processes, as documented in other brain regions^56,57^. Further, cell types involved in social behavior and homeostatic control appear more inter-connected, at least via neuropeptides, at P10 than at P65, consistent with a higher reliance on social interactions for homeostatic needs early in life^58^.

### Development of transcriptional sex differences in POA cell types

The POA and surrounding regions are hotspots of transcriptional sex differences in the brain^29^. Strikingly, we found widespread differences in maturation timing when we examined the maturation trajectories of POA cell types (Fig 2a) in males and females separately (Fig 4a-e and Extended Data Fig 8a). POA cell types were more mature in females than males at the earliest stages of our study (E18 to P4) but became more mature in males than females between P4 and P10 and remained so through P28 (Fig 4c-d and Extended Data Fig 8a). The higher maturation in males at P4-P10 coincides with the timing of transcriptional changes resulting from the perinatal testosterone surge in males^59^. Cell types in females showed more pronounced maturation at P28-P65 (Fig 4c-d), during puberty, suggesting that puberty plays a larger maturational role in females. Beyond maturation timing, sex also affected whether cell types matured gradually or stepwise. Due to earlier and more pronounced maturation at P4-P10 and P18-P28 in males than in females, cell types in males showed more gradual maturation (Class 1), whereas cell types in females matured largely in two steps (Class 3), P10-P18 and P28-P65 (Fig 4a-b, d-e). Intriguingly, these sex differences were present across many cell types, including those with low or undetectable levels of sex hormone receptor expression (Extended Data Fig 8b-c). These observations may result from the lack of sensitivity of snRNA-seq in detecting low Esr1 expression levels but also raise the possibility of indirect effects from sex-hormone-responsive neurons on the maturation of other cell types. Indirect effects of sex hormones could also explain the prolonged maturation of certain cell types involved in social behavior that lack sex hormone receptor enrichment (Fig 2j).

**Fig 4.**
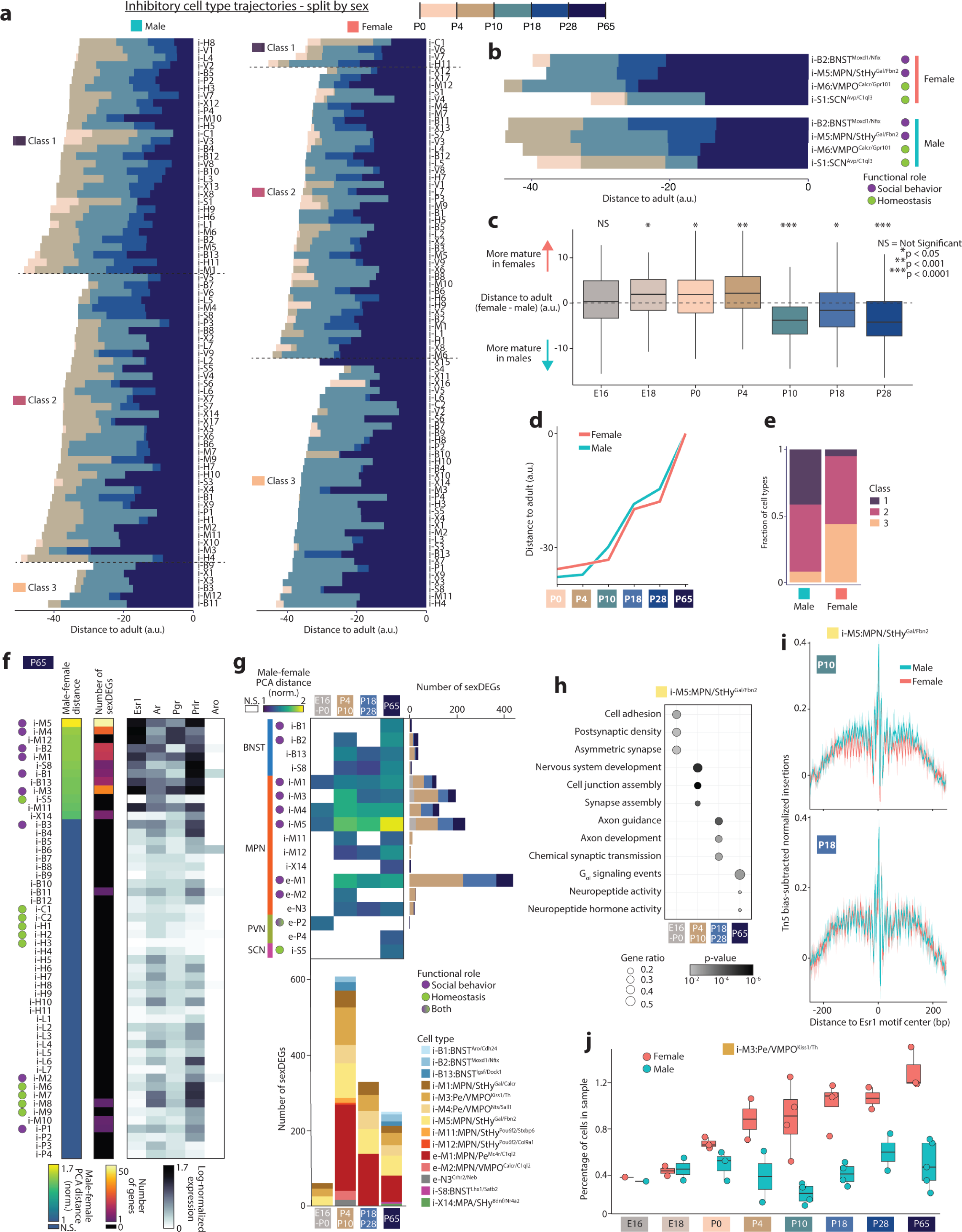
Sex differences in POA maturation. **(a)** Distance trajectories among inhibitory clusters from male (left) or female (right) animals. **(b)** Distance trajectories for four example clusters in females or males. **(c)** Distribution of distance to adult at each age, subtracting male from female distance, per cluster. p-value indicates results from two-sided t-test, male versus female distances, across all cell types. **(d)** Distance trajectories separated by sex averaged across all clusters. **(e)** Fractional class distribution of cluster distance trajectories. **(f)** At P65, sex differences in inhibitory clusters measured by (left) the distance between male and female sample centroids in PCA space (normalized to intra-sex distance) or (middle) number of sexDEGs detected. Right, averaged sex hormone receptor expression. N.S. = not significant. **(g)** All clusters that show significant sex differences in at least one age group according to male-female PCA distance (upper left). Histograms show number of sexDEGs among these clusters per age. N = 6-7 female and 6 male samples at each age, excepting P65 with 3 female and 5 male samples. N.S. = not significant. **(h)** Top GO terms for sexDEGs unique to specific ages in i-M5:MPN/StHy^Gal/Fbn2^. **(i)** Bias-corrected ATAC-footprints centered around the Esr1 motif binding site, quantified separately for male and female i-M5:MPN/StHy^Gal/Fbn2^ cells at P10 (upper) and P18 (lower). **(j)** Percentage of cells identified as i-M3:Pe/VMPO^Kiss1/Th^ in each sex at each age. Each point represents a single sample (3 POAs pooled).

Sex differences in gene expression are concentrated in MPN and periventricular POA (Pe), as well as BNST, emerge due to surges in circulating sex hormones at birth and puberty, and depend largely on expression of the estrogen receptor Esr1^27,29,30,59^. However, the relative contribution of birth and puberty in establishing these sex differences remains unclear, particularly at cell type resolution. Do some cell types show sex differences only following birth, and others only upon puberty? Do perinatal sex differences affect different neuronal functions than post-pubertal ones? To address these questions, we asked which cell types showed sex differences in adults (P65) using two measures: (1) distance between male and female cells in high-dimensional PCA space^41^ and (2) number of differentially expressed genes between males and females^59^ (sexDEGs). These measures gave similar results, converging on a set of 15 out of 147 cell types showing significant sex differences in gene expression (Fig 4f and Extended Data Fig 8b-c). These cell types were enriched in sex hormone receptor expression, particularly Esr1 (Fig 4f and Extended Data Fig 8b-c), and also showed Esr1 motif enrichment in ATAC-seq data relative to other MPN cell types (Extended Data Fig 8d). Many of these cell types have previously been shown to display sex differences in gene expression or function^1,27,28,59^ (Extended Data Table 1).

We next examined earlier ages using the same two methods. In total, 17 cell types showed sex differences at one or more ages (Fig 4g). 14 of these 17 cell types are located in MPN/Pe and BNST, and 7 of these 14 have previously been implicated in sex-differential social behaviors such as mating, parenting, or social drive (Fig 4g). How does birth and puberty affect sex differences among these cell types? At E16-P0, 4 cell types showed significant sex differences, and 61 sexDEGs were found, compared to 12 cell types and 608 sexDEGs at P4-P10 (Fig 4g). At P65, 15 cell types showed sex differences, and 292 sexDEGs were found, 141 of which were also found at earlier ages (Fig 4g). Altogether, these data reveal a dramatic increase in sex differences following birth (8 additional cell types and 547 additional sexDEGs), and a much smaller change in sex differences following puberty (3 additional cell types and 151 sexDEGs gained but 316 lost). Remarkably, many genes showed sex differences in gene expression at P4-P10 but not later ages (Extended Data Fig 8e).

The two cell types that showed the highest degree of sex difference were e-M1:MPN/Pe^Mc4r/C1ql2^ and i-M5:MPN/StHy^Gal/Fbn2^, both of which were recently shown to play sex-specific roles in social behavior^1,6^. A gene ontology analysis of i-M5:MPN/StHy^Gal/Fbn2^ sexDEGs indicated age-specific enrichment in several neuronal and developmental processes (Fig 4h). SexDEGs unique to P4-P10 were enriched in genes related to nervous system development and synapse assembly. SexDEGs unique to P18-P28 were enriched in genes involved in synaptic transmission, as well as axon development and guidance. At P65, sexDEGs were no longer enriched in genes relating to developmental processes but were instead enriched in genes related to neuropeptide and hormone signaling. In e-M1:MPN/Pe^Mc4r/C1ql2^, a similar progression from sexDEGs related to neurodevelopment early in life to specific neuronal functions later in life was observed, albeit with distinct functions such as GABAergic synapses or Trkb and protein kinase C signaling (Extended Data Fig 8e-f). These data indicate that while the perinatal period is the key stage for establishing sex differences in these two cell types, sex-specific expression changes continue at later ages in a cell-type specific manner to affect neuronal development and function.

To determine how sex hormone signaling might underly dynamic sex differences in gene expression, we examined Esr1 motifs in our chromatin accessibility dataset. Transcription factor footprinting analysis supported higher accessibility at Esr1 motifs in males compared to females in i-M5:MPN/StHy^Gal/Fbn2^ neurons at P0 and P10, but not at P18 (Fig 4i and Extended Data Fig 8g). This suggests that the perinatal testosterone surge can exert sex-specific effects via Esr1 through P10, but that Esr1 function is no longer sex-specific by P18, which may partly explain dynamic changes in gene expression across age (Fig 4h).

The cell type i-M3:Pe/VMPO^Kiss1/Th^ was the only population to show a replicable sex difference in cell number (Fig 4j), with a 3:1 female bias as previously shown^1,28^. Sex differences in transcriptomic cell type number have also been reported in BNST^1,59^, which was not apparent in our dataset, perhaps due to variability in BNST dissection. Differences in BNST cell number arise perinatally via estrogen-dependent protection from apoptosis in males but not females^27^. In contrast, how and when female-biased cell number differences emerge in i-M3:Pe/VMPO^Kiss1/Th^ is not known. Our data indicate that, similar to BNST cell types, the sex bias in cell number in POA emerges perinatally (Fig 4j). While i-M3:Pe/VMPO^Kiss1/Th^ showed numerous sexDEGs before and after birth, we did not observe sex differences in genes related to apoptotic pathways. Finally, although microglia and prostaglandin signaling have been implicated in the emergence of perinatal sex differences^60^, low cell numbers prevented us from assessing sex differences in these cells, and we did not find sexDEGs associated with prostaglandin signaling.

In conclusion, our dataset reveals widespread sex differences in the mode and timing of POA cell type maturation. We uncover transcriptional programs underlying neuronal development and function that exhibit strong sex biases in an age-restricted manner and in a small number of key cell types, with the largest effects seen in cell types known to be involved in social behavior. These findings illustrate how sex hormones play a global yet cell-type-specific role in shaping POA development.

### Sensory inputs affect POA cell type maturation

Proper maturation of brain regions involved in sensory processing relies on spontaneous and sensory-evoked neuronal activity^7–10^. Whether sensory inputs also affect the maturation of brain regions involved in homeostatic control or social behavior, such as the POA, is unclear^17^. To address this question, we focused on sensory modalities that are crucial for POA function: somatosensation, including thermoregulation^4,54^ and social touch^6,61^, and chemosensation, including the main olfactory system, essential for mating and aggression^62,63^, and the vomeronasal system, required for pheromone detection and the sex-specificity of social behaviors^18,64–66^. We performed single-nucleus sequencing on POAs from mutants affecting each of these sensory modalities, focusing on P10, P18, and P65 as maturational changes are concentrated at those ages (Fig 2). For each mutant, we asked which cell types differed from controls using (1) distance between mutant and control cells in high-dimensional PCA space (as was used to assess sex differences in Fig 4) and (2) a random forest classifier.

Mutants with conditional knockouts of Piezo2 or Gabrb3, which lead to touch hypo- or hyper-sensitivity as early as before birth^67,68^, showed no POA cell types with significant differences compared to littermate controls (Fig 5a for inhibitory cell types and Extended Data Fig 9a for excitatory cell types; both in Extended Data Fig 9b). In Trpm8 mutants, which are strongly impaired in cold sensation^69^, four cell types showed significant differences at P10, including e-X1:POA^Bmp7/C1ql1^ which partially localizes to the thermoregulatory nuclei AvPe/MnPO, but no cell types were affected at P18. Altogether, these data indicate a minimal role for somatosensory input in the transcriptomic maturation of POA cell types.

**Fig 5.**
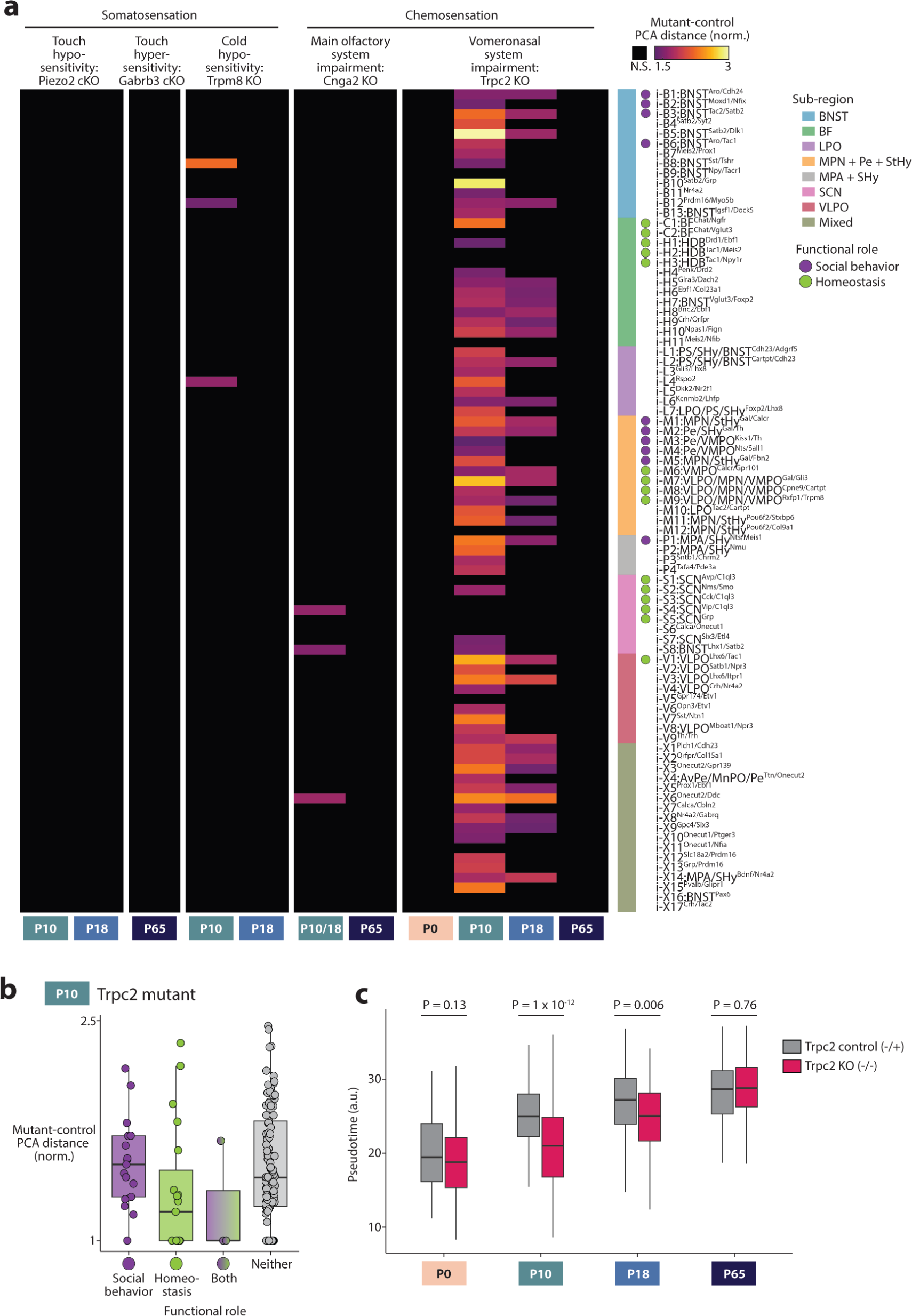
Sensory signals affecting POA maturation. **(a)** Distance between mutant and control sample centroids in PCA space (normalized to intra-genotype distance) for inhibitory clusters. Control samples included both C57BL6/J and control littermates. Sample numbers, left to right columns, mutant/control: 2/8, 4/10, 2/10, 2/8, 2/8, 2/6, 2/7, 2/7, 2/8, 2/8, 2/7. **(b)** Mutant-control PCA distance for all clusters in Trpc2 P10 experiments, split according to functional role. **(c)** Pseudotime mapping of Trpc2 mutant (-/-) and control (-/+ littermates) across age. Pseudotime is averaged within a cluster, boxplot shows distribution across all cluster averages. p-value indicates results from two-sided t-test, mutant versus control cell types.

We next examined mutants with loss of main olfactory (Cnga2 KO)^70^ or vomeronasal (Trpc2 KO)^64^ function. Seven POA cell types showed small but significant differences in Cnga2 mutants compared to controls at P10 and P18, whereas none were affected at P65 (Fig 5a and Extended Data Fig 9a-b). Surprisingly, the majority of cell types in Trpc2 mutants showed a significant difference between mutants and controls at P10, with many cell types showing a 2-3 fold higher distance between versus within genotypes (Fig 5a, Extended Data Fig 9a-b). These effects were absent at P0, largest at P10, smaller at P18, and small but detectable at P65 (Fig 5a, Extended Data Fig 9a-b), indicating a mostly transient effect that coincides with the timing of major maturational changes in POA (Fig 1-4). Because Trpc2 expression is restricted to the vomeronasal organ^71^we ascribe this result to impaired vomeronasal function rather than effects in other brain areas.

Pheromonal information from vomeronasal sensory neurons reaches POA via accessory olfactory bulb projections to BNST and medial amygdala^72^. In Trpc2 mutants at P10, cell types in BNST were particularly affected, as were cell types in various POA sub-regions, whereas cell types outside the POA in regions with little involvement in social behavior, such as SCN or BF, were less affected (Fig 5a and Extended Data Fig 9c). Cell types involved in social behavior were more affected than cell types involved in homeostatic control (Fig 5b), a result consistent with the importance of vomeronasal signaling in mouse social behavior control and the major projections from the vomeronasal system to the preoptic region^72^. Surprisingly, some effects were also seen on cell types not expected to rely on vomeronasal signaling, for example VLPO^Tac1^ neurons involved in sleep and VMPO^Calcr^ neurons involved in fever generation and sickness behavior, suggesting either unexpected vomeronasal connectivity or indirect developmental effects from neighboring cell types. To validate these results and assess the specificity of the widespread effects observed on POA cell types, we asked whether maturation in other brain areas was affected. For this purpose, we collected Trpc2 mutant and control littermate brains at P10 and dissected both POA and visual cortex from the same brains. As in our initial experiment, the majority of POA cell types were significantly different in Trpc2 mutants versus control littermates, whereas visual cortex cell types were largely unaffected (Extended Data Fig 9d-e).

POA cell types appear different in Trpc2 mutants mostly at P10 and to a lesser extent P18 (Fig 5a and Extended Data Fig 9a-b). This could indicate improper developmental timing, such as transiently accelerated or delayed maturation, or an alternative developmental route that later converges toward the wild-type adult state. To address this issue, we quantified a pseudotime trajectory^73^ for each cell type using our wild-type dataset across all eight ages, which gives a maturation score to each cell. We then projected Trpc2 mutant and littermate control cells onto the wild-type pseudotime axis to determine their maturation states (Fig 5c). As expected, both Trpc2 control and mutant cell types showed increasing pseudotime with age. However, at P10 and P18 but not P0 or P65, Trpc2 mutant cell types showed significantly lower pseudotime values than controls (Fig 5c), consistent with a transient developmental delay. Cell types in Trpc2 mutants at P10 resembled those in littermate controls at P0 (Fig 5c, P = 0.99). In summary, maturation of POA cell types appears to occur largely independently from somatosensory or main olfactory input, yet, for the majority of POA cell types, particularly, though not exclusively those involved in social behavior, requires vomeronasal input for proper developmental timing.

## Discussion

In this study, we explored transcriptional and chromatin dynamics underlying POA cell type maturation, and uncovered key determinants of developmental trajectories, including the specific preoptic subregion in which a cell type resides, its ascribed function, the sex of the animal, and the presence of functional input from specific sensory modalities. POA cell types appear already diversified at E16, soon after neurogenesis, which differs from developmental timelines in cortex^74,75^ or sensory systems^76,77^ where subtype diversification continues well beyond neurogenesis and into postnatal life. A wide diversity of neuronal types was also observed in the Arcuate nucleus of the hypothalamus at E15.5^78^, although these embryonic neuronal types were not as clearly related to adult cell types as we find in POA.

Following this early diversification, POA cell types undergo several key maturational steps. Shortly after birth between P0 and P10, sub-regional identity strengthens (Fig 1), signaling networks are established (Fig 3), and vomeronasal input ensures proper maturational timing (Fig 5). In a subsequent step, signaling networks are reconfigured, and cell types become increasingly adult-like, with two critical maturation stages identified as P10-P18 and P28-P65 (Fig 2). Intriguingly, P10-P18 just precedes mouse pup weaning (P18-P28) and has been identified in various brain regions as a key time point for brain maturation both molecularly and physiologically^79–81^. P28-P65 encompasses puberty (P25-P42), and cell types involved in social behavior show extensive maturation at this age in particular. We find that sex affects maturation at all time points, including the perinatal emergence of sex differences in Esr1+ cell types, widespread differential maturation trajectories, and dynamic cell-type-specific sex differences in gene expression upon puberty. Overall, our results indicate that the precise timeline of maturation is highly cell-type-specific and occurs within different (early or late) time windows and in a gradual or stepwise manner, according to the cell sub-regional identity, behavioral function, and sex.

Finally, we examined POA cell type maturation in mutants affecting various sensory modalities and found little effect of somatosensory or main olfactory impairment. Clear caveats of these results are the lack of sensitivity of snRNA-seq approaches to lowly-expressed genes, the poor representation of some cell types in our dataset, the inclusion of only key postnatal time points, and partial removal of sensory inputs (e.g. remaining touch signals above the neck in Piezo2 cKO animals or heat sensation in TrpM8 knockouts). By contrast, in animals with loss of vomeronasal function, we found a major effect on maturational timing on the majority of cell types. A role for vomeronasal input in POA maturation suggests that input-dependent plasticity during development occurs beyond sensory and cognitive systems, in brain regions classically assumed to be genetically hard-wired and to show little plasticity^17,26,82^. How and why POA maturation depends so strongly on vomeronasal input is an interesting question that may be addressed in future work. In Trpc2 mutants, vomeronasal sensory neurons fail to respond to pheromones^64^, but display normal spontaneous neuronal activity, which in other sensory systems is a key neurodevelopmental factor. Whether late embryos and newborns sense vomeronasal cues has not been explored, nor is it known whether vomeronasal signals may affect POA neuronal activity in early life. Vomeronasal inputs reach some POA cell types (e.g. Gal+ neurons^19^) and not others (e.g. Gnrh+ neurons^62^). Given the widespread developmental effects of lack of VNO sensory activity on many POA cell types, one might hypothesize that cell types receiving direct vomeronasal input may be initially affected, and in turn fail to release key signaling molecules, thus indirectly affecting the maturation of neighboring cell types. Our NeuronChat analysis identified several POA clusters, including those affected in Trpc2 mutants and involved in social behavior, that signal via distinct neuropeptides at P10 versus P65 and may thus play key maturational roles. In adults, we see smaller differences between Trpc2 mutant and control POA cell types, which may or may not be functionally relevant for adult function. Importantly, even a transient transcriptional difference during development can lead to permanent and functionally relevant differences in connectivity and other neuronal features, as shown in other systems^83^. Based on the major effect of vomeronasal input on the developmental timing of POA cell types, we propose that ∼P10 may represent a critical period for POA circuit establishment and sensitivity to vomeronasal input.

Altogether, our work shows that sensory input, sex, sub-region localization, and function impinge upon POA cell type development. This points to a developmental process that is surprisingly sensitive to external influences, such as vomeronasal input, sex hormone secretion, and subregion-specific local signaling. How might adult homeostatic or social behavior function affect changes during development? This could result from a combination of genetically pre-programmed (e.g. lineage-based) effects and environmental influences, such as vomeronasal and sex hormone inputs as well as effects associated with age-dependent life changes such as the onset of thermogenesis and independent feeding. The time-restricted sensitivity of POA development to vomeronasal input is highly reminiscent of critical periods of sensory development in which activity-dependent mechanisms operating within certain time-windows affect aspects of development and later function. Altogether, the sensitivity of POA development to external inputs suggests underlying mechanisms by which early life experience may lead to long-lasting effects on social behavior or homeostatic control. Thus, as proposed by Nikolaas Tinbergen in his Four Questions^13^, studies of development may help address long-standing questions on the origin of instinctive behaviors.

## Extended Data Figure Legends

**Extended Data Fig 1.**
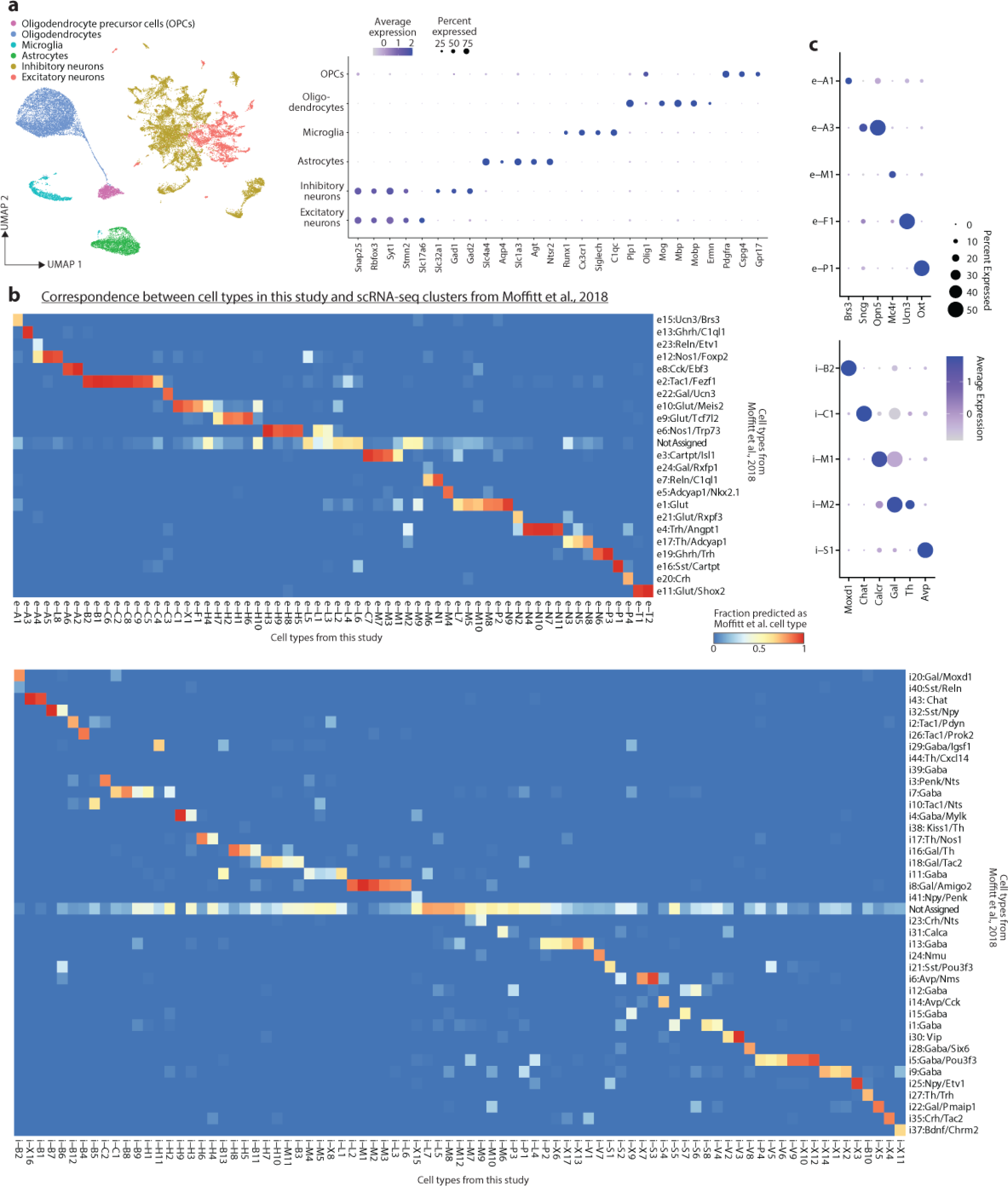
Identifying major cell classes and mapping cell types to reference. **(a)** UMAP colored by cell class (excitatory/inhibitory neurons, oligodendrocytes, astrocytes, microglia) and associated Dotplot with gene markers to verify class IDs. **(b)** Mapping of cell types in this study (columns) to scRNA-seq cell types in the POA reference atlas (Moffitt et al., 2018), for excitatory (upper) and inhibitory (lower) neurons. Canonical correlation analysis was used to generate a prediction score, which was then thresholded (Methods) to give predicted cell identity or NA for cells that did not pass threshold for any cell type. Color represents the fraction of cells in each cell type in this study that were predicted to belong to each reference atlas cell type. **(c)** Dotplot showing gene markers for previously studied cell types (see Fig 1b for cell type function and full name).

**Extended Data Fig 2.**
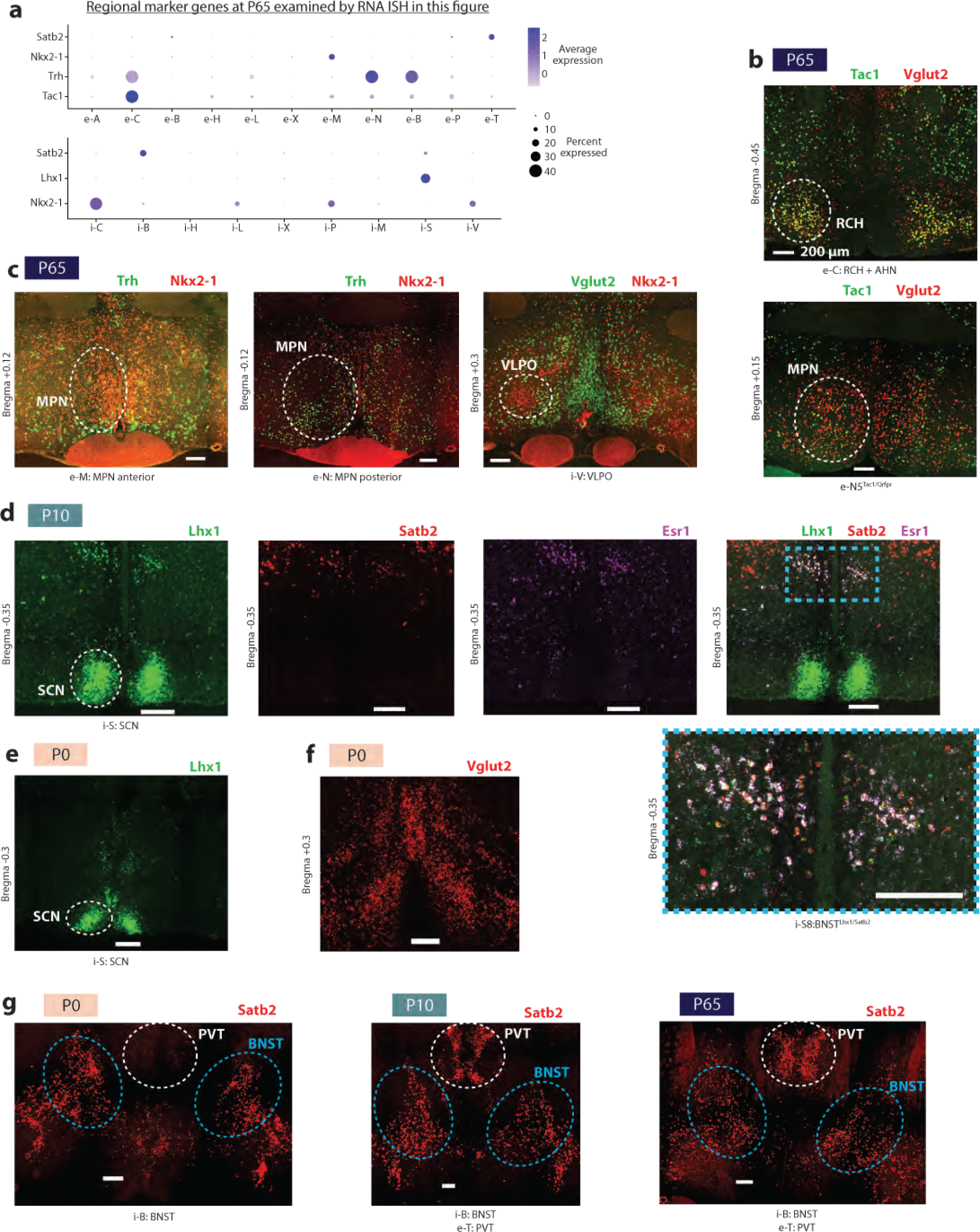
RNA in situ hybridization confirmation of cell type localization and changes across age. **(a)** Regional markers at P65, for genes stained using ISH in subsequent panels of this figure. **(b)** Upper: Tac1+ Vglut2+ neurons residing in RCH. Lower: A small population of Tac1+ Vglut2+ neurons resides in MPN, corresponding to e-N5^Tac1/Qrfpr^ **(c)** Trh and Nkx2-1 differentiate anterior and posterior MPN excitatory neurons. Left: Nkx2-1+ and Trh-cells reside in anterior MPN. Many of these neurons are Vglut2+ (data not shown, but co-stained). Middle: Trh+ and Nkx2-1-cells reside in posterior MPN. These neurons are also Vglut2+ (data not shown, but co-stained). Right: a population of Nkx2-1+ and Vglut2-neurons demarcates VLPO. **(d)** Left: Lhx1 is a marker for SCN neurons at P10 (and other ages). Other panels identify a population of BNST neurons, termed i-S8:BNST^Lhx1/Satb2^, that co-expresses Lhx1, Satb2, and Esr1. **(e)** Lhx1 is a marker for SCN neurons also at P0. **(f)** Vglut2 staining shows the inverted-Y structure of AvPe/MnPO at P0, as in adults (panel **c**, right) **(g)** Satb2 is a marker for BNST at P0, P10, and P65, but for PVN only from P10 onwards, not at P65.

**Extended Data Fig 3.**
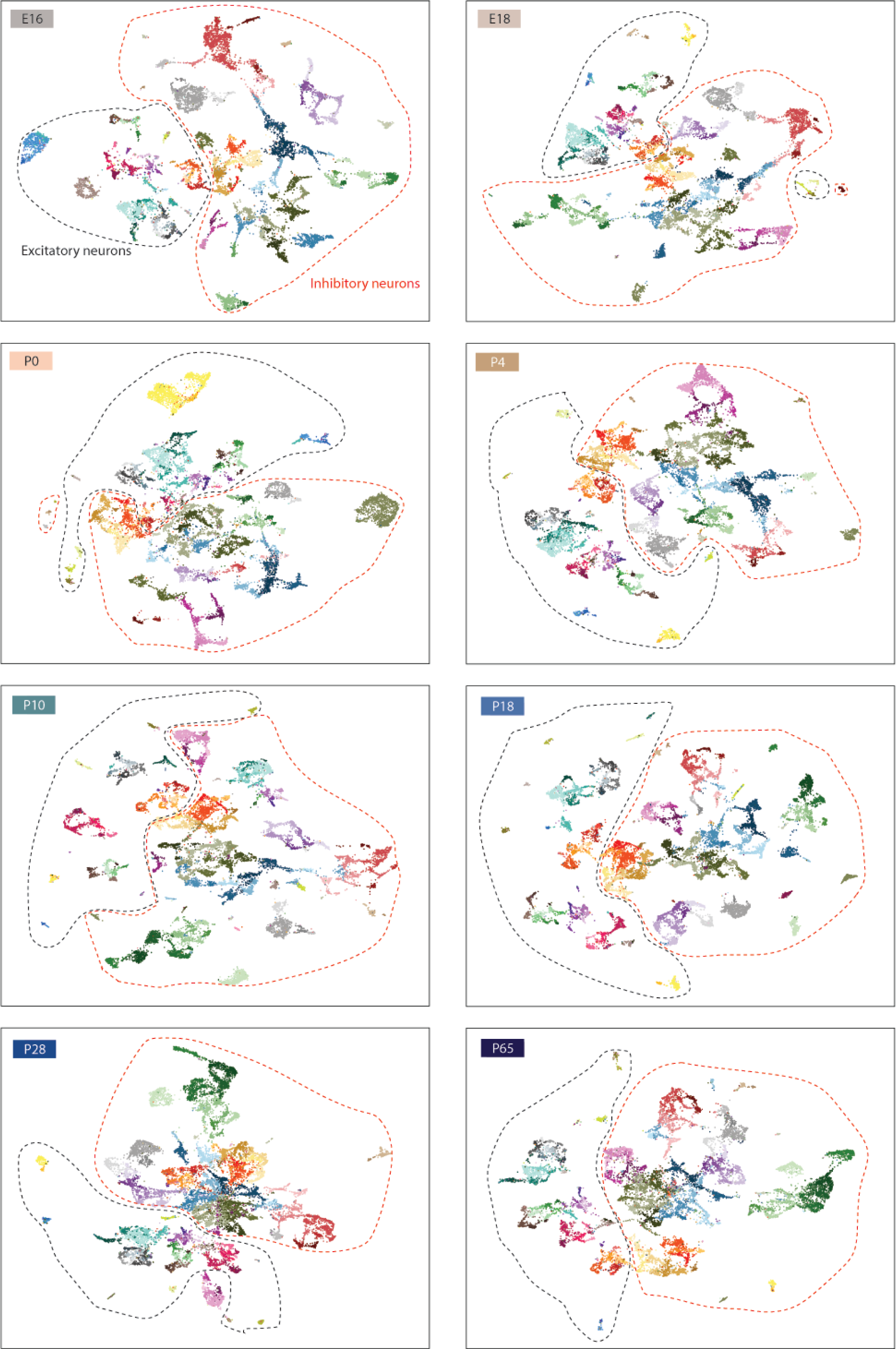
UMAPS with cell type IDs across all ages. Excitatory and inhibitory neurons plotted together. Dotted lines encircle excitatory (black) or inhibitory (red) neurons. See key in Fig 1c for cell type color labels.

**Extended Data Fig 4.**
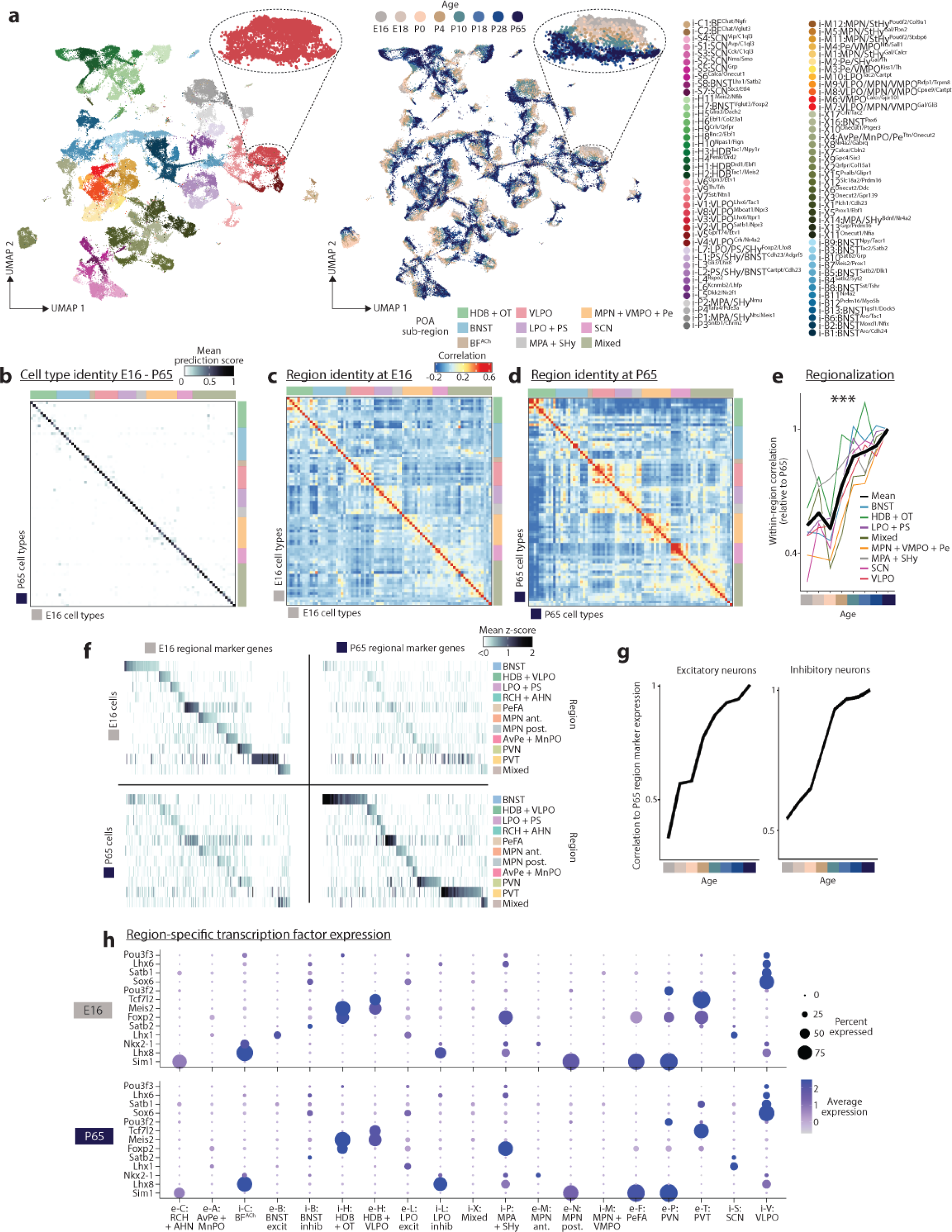
Region-specific gene expression changes with age. **(a)** UMAP across all 8 ages for inhibitory neurons. Left, colored by cluster (key in Fig 1c, upper); right, colored by age. Inset: gradient across age for one cluster. **(b)** Mapping of E16 onto P65 clusters using canonical correlation analysis, across inhibitory neurons. Prediction score is averaged across cells within a cluster. **(c)** Correlation between pseudobulked gene expression of all clusters at E16, across inhibitory neurons. **(d)** As in (c) but at P65. **(e)** Correlation between clusters within a sub-region, at each age, normalized to correlation at P65. Analysis shown for inhibitory neurons (Fig 1 shows excitatory neurons). ***p < 0.0001 for one-way ANOVA, effect of age. **(f)** Marker genes specific to excitatory neuron sub-regions, found at either E16 but not P65 (left) or P65 but not E16 (right), plotted for E16 cells (upper) or P65 cells (lower). **(g)** Using regional markers at P65 that are not markers at E16 (genes on right panel of (f)), correlation of regional expression at each age compared to P65. **(h)** Region-specific marker genes at P65 (upper) or E16 (lower).

**Extended Data Fig 5.**
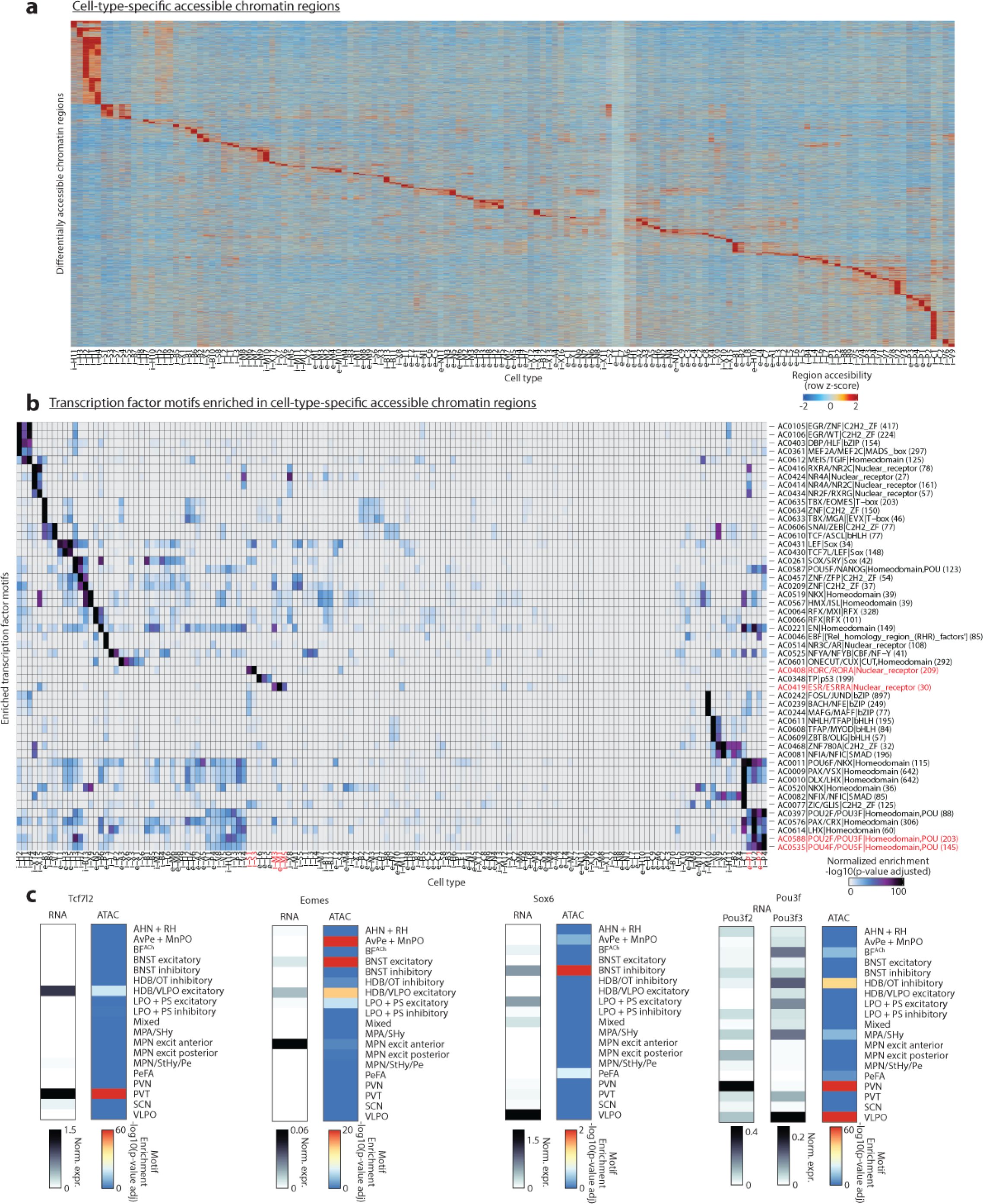
ATAC-seq identifies cell-type- and region-specific transcription factor motifs. **(a)** Differentially accessible regions (DARs) of chromatin specific to each cell type. **(b)** Cell-type-specific transcription factor motif enrichment, identified from cell-type-specific DARs. Motifs and cell types discussed in the main text are highlighted in red. **(c)** Examples of transcription factor motifs in accessible chromatin (ATAC-seq data) enriched amongst anatomical sub-regions that match with region-specific gene expression.

**Extended Data Fig 6.**
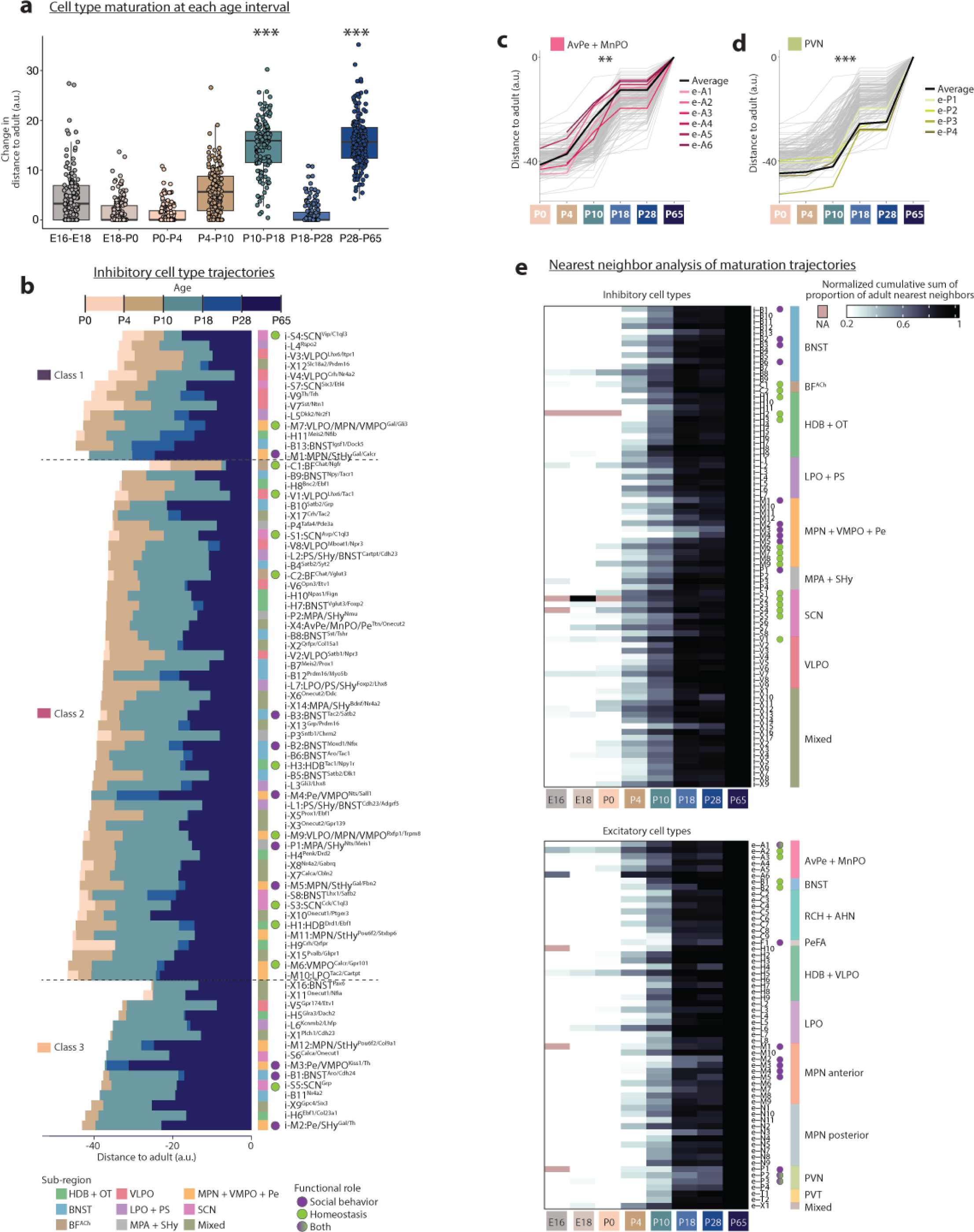
Maturational trajectories of POA cell types. **(a)** Difference between distance to adult at each subsequent pairs of ages. Each point represent a single cell type. ***p < 0.0001 for one-way ANOVA with post-hoc Tukey’s test comparing those P10-P18 and P28-P65 to all other ages (except each other). **(b)** Distance trajectories of inhibitory clusters. **(c)** Distance trajectories for AvPe + MnPO clusters. **p = 0.000198 for two-way ANOVA, effect of cell type subset. **(d)** Distance trajectories for PVN clusters. ***p < 0.0001 for two-way ANOVA, effect of cell type subset. **(e)** Heatmaps of scaled cumulative sum of proportion of nearest neighbors that are nuclei from adult (P65). Upper: inhibitory cell types. Lower: excitatory cell types.

**Extended Data Fig 7.**
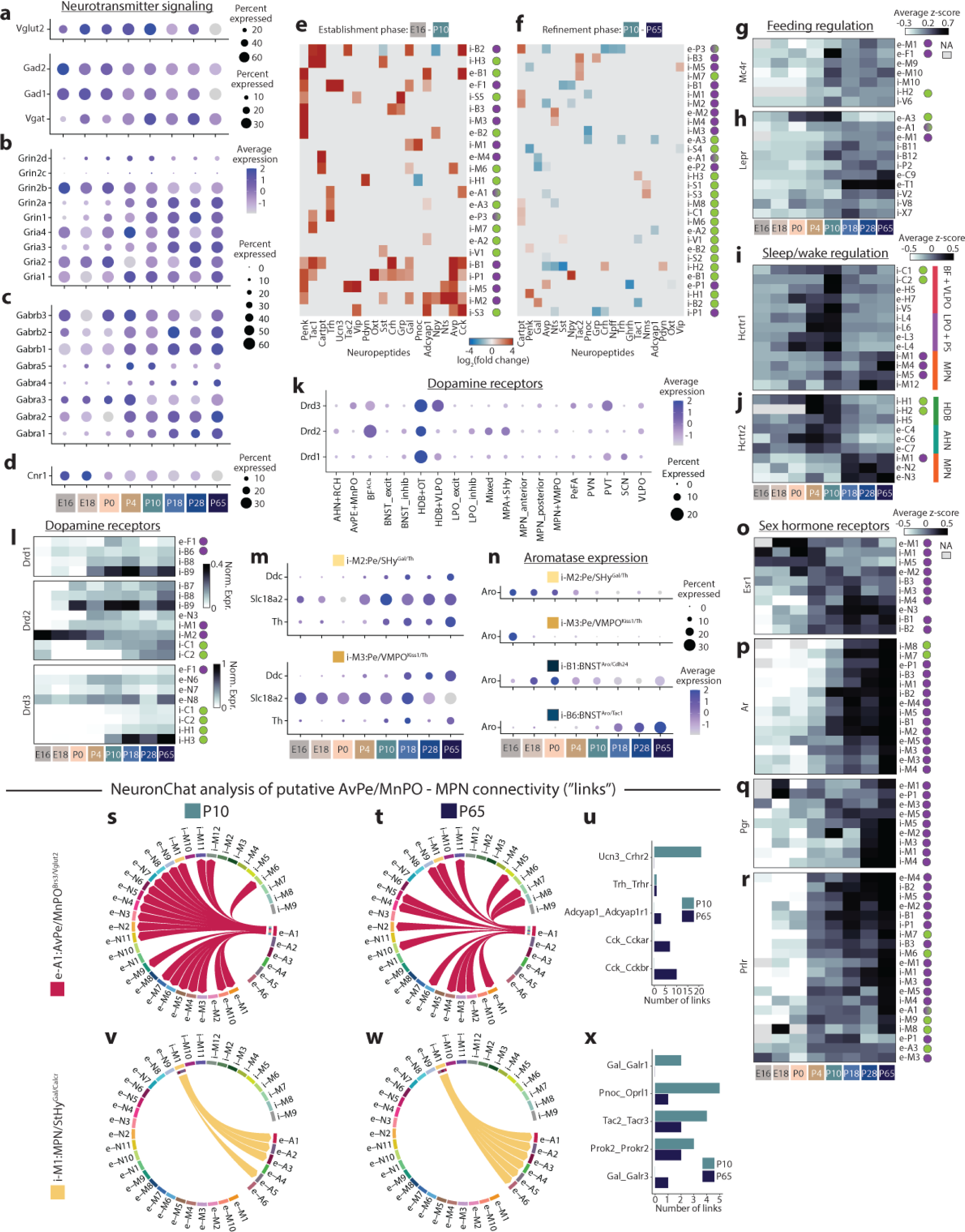
Developmental changes in neuronal signaling gene expression. **(a-d)** Dotplots showing expression across all neurons, separated by age, for glutamate and GABA synthesis and packaging enzymes (a), selected glutamate receptors (b), selected GABA receptors (c), or endocannabinoid receptor Cnr1 (d). **(e-f)** Heatmap showing significant increases (red) or decreases (blue) in expression of neuropeptide genes between E16 and P10 (e) or P10 and P65 (f). Gray indicates no statistical significance. **(g-j)** Heatmaps showing expression of selected genes involved in feeding behavior (g, Mc4r; h, Lepr) or sleep/wake behavior (i, Hcrtr1; j, Hcrtr2) across cell types in which those genes are enriched. Expression is z-scored within a cell type and then averaged across cells at each age. **(k)** Dotplot showing expression of dopamine receptors Drd1, Drd2, and Drd3 at P65, split by sub-region. **(l)** Heatmaps showing expression of Drd1, Drd2, and Drd3 across cell types in which these genes are enriched. Expression is log-normalized. **(m)** Dotplots showing expression of genes involved in dopamine synthesis and packaging in the two cell types that express all three of these genes, across age. **(n)** Dotplots showing change across age of aromatase expression in the four aromatase-enriched cell types. **(o-r)** Heatmaps showing expression of sex hormone receptors (o, Esr1; p, Ar; q, Pgr; r, Prlr) across cell types in which these genes are enriched and which have some functional annotations for social or homeostatic behaviors. **(s-x)** NeuronChat analysis of putative output connections from e-A1 (s-u) or i-M1 (v-x). Output cell types are shown at P10 (s,v) or P65 (t,w), with output links compared at these two ages in (u, x).

**Extended Data Fig 8.**
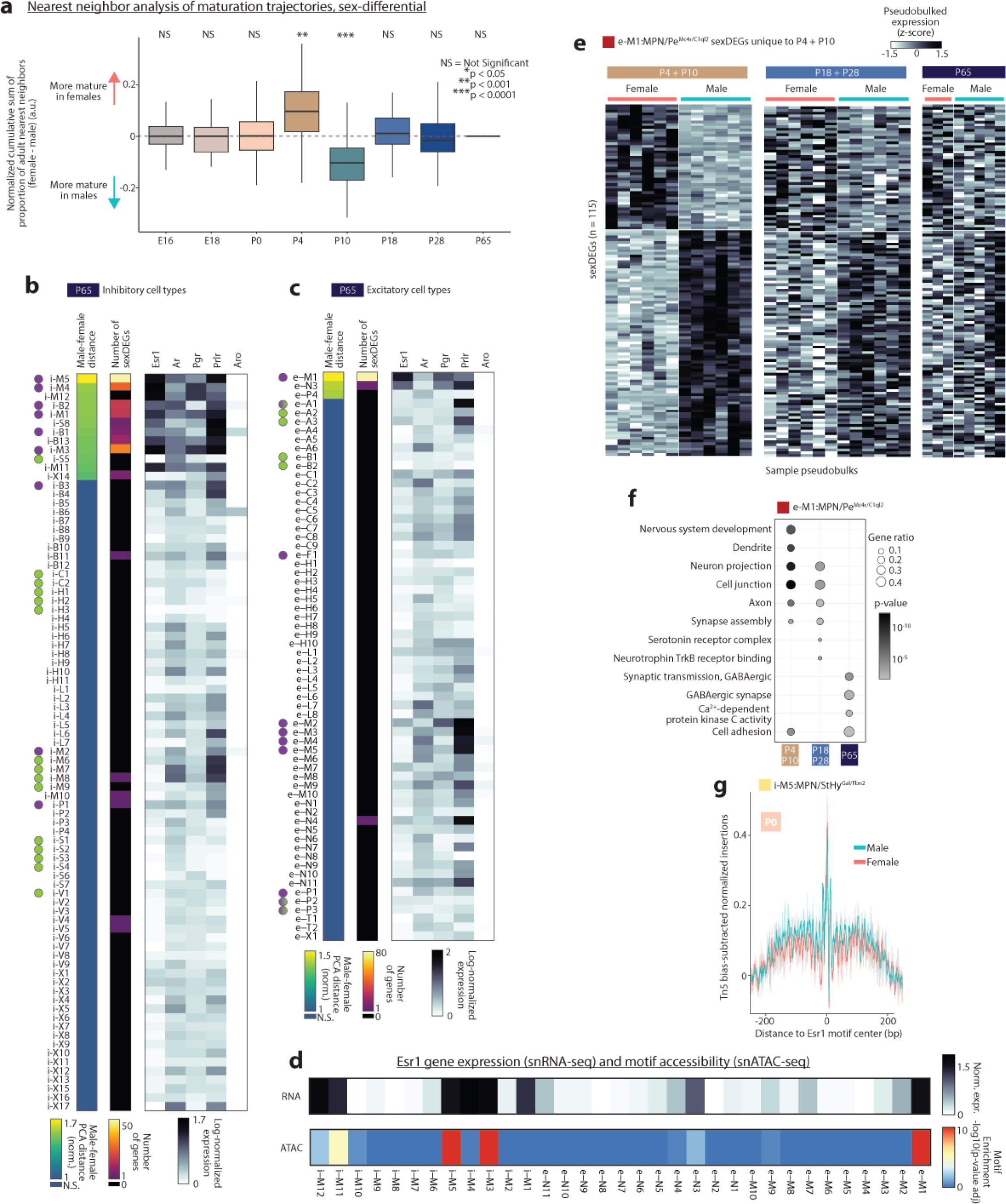
Age- and cell-type-specific sex differences in POA gene expression. **(a)** Difference between male and female proportion of adult nearest neighbors at each age. p-value indicates results from two-sided t-test, male versus female distances across all cell types. **(b-c)** At P65, sex differences in inhibitory (b) or excitatory (c) clusters measured by distance between male and female sample centroids in PCA space (normalized to intra-sex distance; left column) or number of sexDEGs detected (middle column). Grayscale matrix on the right indicates sex hormone receptor expression. **(d)** Esr1 gene expression (upper) and cell-type-specific motif enrichment in accessible chromatin (lower) amongst MPN cell types. **(e)** Gene expression values for 115 sexDEGs detected at P4-P10 but not other ages in e-M1, pseudobulked across cells within samples (each column represents a single sample). Control samples from mutant/control experiments included to increase sample number. **(f)** Top GO terms for sexDEGs found only at specific ages in e-M1. **(g)** Bias-corrected ATAC-footprints centered around the Esr1 motif binding site, quantified separately for male and female i-M5:MPN/StHy^Gal/Fbn2^ cells at P0.

**Extended Data Fig 9.**
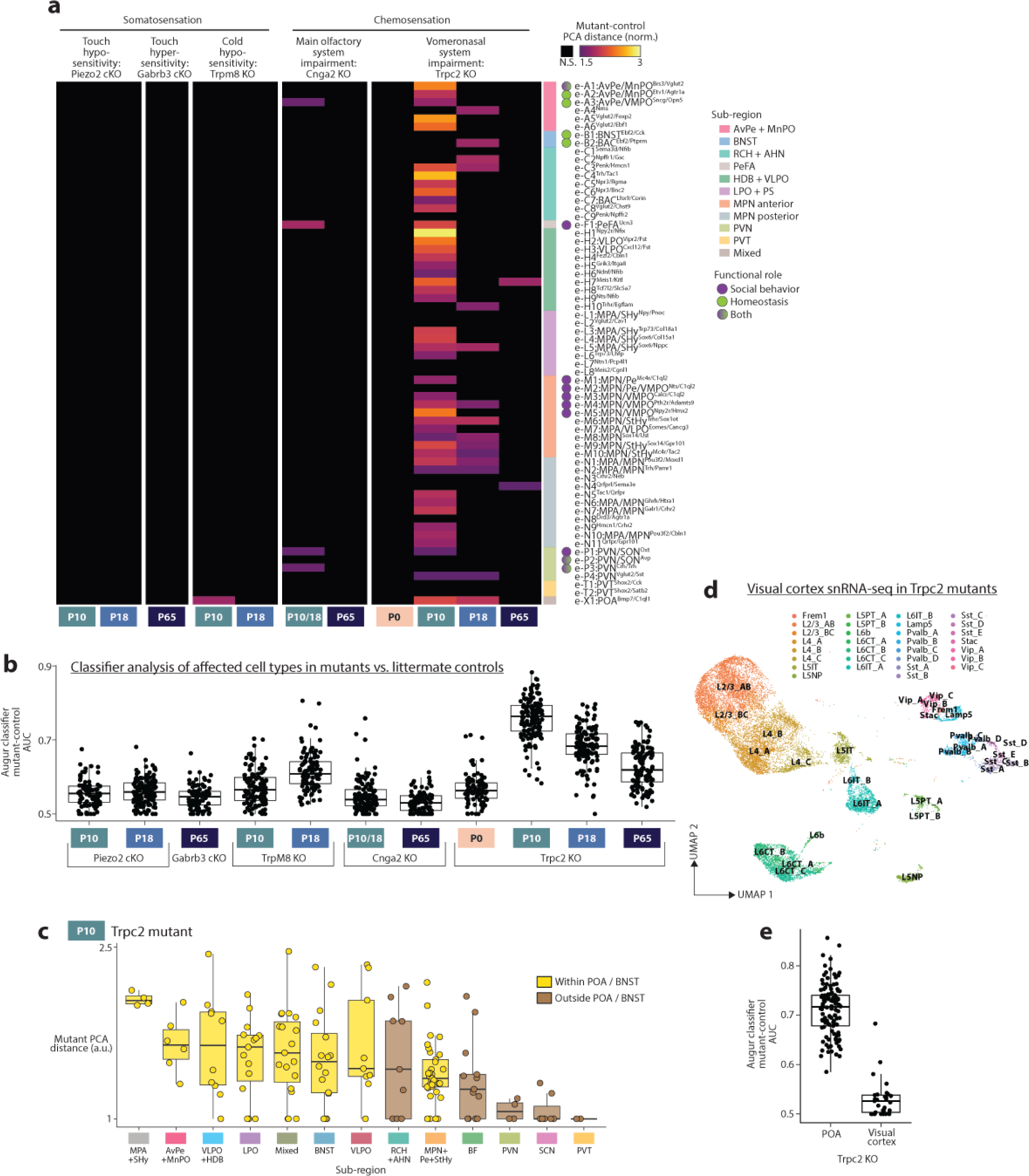
Effects of impaired sensory input on POA development. **(a)** Distance between mutant and control sample centroids in PCA space (normalized to intra-genotype distance) for inhibitory clusters. Control samples included both C57BL6/J and control littermates. Sample numbers, left to right columns, mutant/control: 2/8, 4/10, 2/10, 2/8, 2/8, 2/6, 2/7, 2/7, 2/8, 2/8, 2/7. **(b)** Random forest classifier approach (Augur) to identify cell types that show differences between mutant and control samples. Each point represents a single cell type. **(c)** Mutant-control PCA distance for all clusters in Trpc2 P10 experiments, split according to sub-region, and ordered from most to least affected. **(d)** UMAP showing identification of visual cortex neuronal subtypes in Trpc2 mutants and controls (pooled) at P10. **(e)** Random forest classifier (Augur) for replication of Trpc2 P10 POA (left) and visual cortex (right) from the same brains.

**Extended Data Table 1 | Correspondence between cell types identified here and those identified in Moffitt et al. and previous functional studies.**

**Supplementary Table 1 | Transcriptomic sub-regional groups and associated POA sub-regions.** Each sub-regional transcriptomic group contains neurons belonging to one or more POA sub-regions. See Extended Data Table 1 for information on MERFISH cell type matches that inform this table, along with Extended Data Figure 2.

**Supplementary Table 2 | Signaling gene lists used in Fig 3.**

**Supplementary Table 3 | Additional neuropeptide-receptor links added to the NeuronChat database.**

## Methods

### Animals

Mice were maintained on a 12 h:12 h dark–light cycle with access to food and water *ad libitum*. The temperature was maintained at 22 °C and the humidity was controlled at 30–70%. All experiments were performed in accordance with NIH guidelines and approved by the Harvard University Institutional Animal Care and Use Committee. Except for experiments on mutant strains affecting sensory input, C57BL/6J mice were used for all sequencing experiments. Mothers of all C57BL/6J experiments subjects were placed in a fresh cage when embryos were ∼E12. For samples collected at P28 and P65, animals were separated from parents and opposite-sex siblings at P21, and group-housed, and kept separated from the opposite sex until dissection. C57BL/6J brains were collected during the first three hours of the dark cycle. Cnga2, Trpc2, and Trpm8 mutants were treated the same, whereas Piezo2 cKO and Gabrb3 cKO animals and littermate controls were raised in another facility (Harvard Medical School) and thus brains were collected over a wider time window (2-7 hours into the start of the light phase of 12h:12h light cycle). All mutant experiments were performed with equal numbers of male and female samples, excepting Cnga2 KOs where only males were used due to Cnga2’s location on the X chromosome and impairments of Cnga2 mutants in suckling and mating. Cnga2 KO animals, always males, were generated by crossing heterozygous mothers to FVB fathers leading to hemizygous male progeny. Piezo2 cKO animals were generated using Cdx2-Cre, which drives recombination in neurons (including touch sensory neurons) and some peripheral tissues below cervical level C2, and either Piezo2^fl^/Piezo2^fl^ or Piezo2^fl^/Piezo2^-^. Piezo2 cKO brains were collected at P10 or P18/P19, and exhibit motor defects, thus were frequently provisioned with extra food and gel packs placed on the cage floor; due to these defects, we did not raise animals beyond P19 for experiments in adults. We also did not profile TrpM8 KO animals as adults, because thermoregulation maturation occurs mostly before P18. Gabrb3 cKO animals were generated using Advillin-Cre, which drives recombination in all DRG and trigeminal sensory neurons, and heterozygous Gabrb3^fl/+^ animals, which is sufficient to cause touch hypersensitivity^68^. Gabrb3 cKO brains were collected at P60-P70. Mutant and transgenic strains used in this study were previously published and include (with Jackson Laboratories numbers where applicable): Trpc2 knockout mice^64^ (021208); Cnga2 knockout mice^70^; Trpm8 knockout mice^84^ (008198); Piezo2^fl^ ^85^ (027720); Piezo2^-^ ^86^; Cdx2-Cre^87^ (009350); Advillin-Cre^88^; Gabrb3^fl^ ^89^. Sample sizes were chosen to match or exceed those in similar experiments from relevant studies.

### Single nuclei sequencing

Brains were dissected (P10+) or whole heads were removed (E16 to P4) and frozen in Optimal Cutting Temperature medium and placed at -80℃. The preoptic area and surrounding regions were dissected on a cryostat at -15℃ using the Paxinos Developing and Adult Brain Atlases^90,91^ to determine locations for dissection based on landmarks. Tissue was collected from the anterior-most portion of MnPO, determined based on the position of the anterior commissure relative to the lateral ventricle, to the posterior-most portion of the POA, determined based on the amount of tissue collected past the start location guided by the atlases. Dissected tissue was placed in a 2-mL tube and kept at -80℃ until the day of library preparation. For library preparation, 4-8 samples were prepared simultaneously, pairing mutant/control and male/female samples on the same day. Dissected tissues from 3 mice (C57BL6/J samples) or 2-3 mice (mutant/control samples) were pooled for each sample. 2 female and 2 male samples were collected at each of the 8 ages, except for E16, where only 1 female and 1 male sample were collected. Nuclei preparation was modified from 10x Genomics demonstrated protocol CG000375. Samples were dounce homogenized in a 3mL Potter-Elvehjem Tissue Grinder in a 4℃ cold room in 0.1% NP40 lysis buffer, passed through a 70-µm filter (MACS Smart-strainer) and centrifuged at 500g for 5 min. All buffers contained 1 U/µl RNAse Inhibitor (Sigma Protector) and centrifugation steps in nuclei preparation was done at 4℃. Nuclei were resuspended in PBS with 2% BSA, then passed through a 20-µm filter (MACS Smart-strainer). 7-AAD was added, and nuclei were separated from debris by FACS. Following centrifugation, nuclei were permeabilized in 1x lysis buffer (10x Genomics CG000375), then centrifuged and resuspended to a concentration of ∼6,000 nuclei per µl, using a Luna Cell Counter. Downstream preparation of sequencing libraries was carried out using the 10x Genomics Multiome Kit. Libraries were sequenced on an Illumina NovaSeq 6000 S4 200 flowcell using instructions provided by 10x Genomics Multiome protocol (GEX: read 1=26bp, i1=10bp, i2=10bp, read 2=90bp; ATAC: read 1=50bp, i1=8bp, i2=24bp, read 2=49bp), to a saturation of at least 70% for each run. Paired-end sequencing with read lengths of 100 nt was performed for all samples.

## Single-nucleus RNA-sequencing analysis

Illumina sequencing reads were aligned to the mouse genome using the 10x Genomics CellRanger ARC pipeline with default parameters. For the C57BL6/J dataset, the mean numbers of UMIs and genes per cell were 5276 UMIs and 2264 genes for all neurons. Raw reads and output are available at: https://data.nemoarchive.org/biccn/grant/u19_huang/dulac/transcriptome/sncell/10x_multiom <u>e/</u>

For initial analysis, we relied on the R package Seurat and standard data analysis practices^92^. We filtered out nuclei with under 400 or over 100,000 UMIs, with under 250 genes, and with over 20% of UMIs belonging to one gene. While rare, nuclei with over 10% mitochondrial or ribosomal UMIs, or over 1% IEG, apoptotic, or red blood cell UMIs, were also filtered out. We defined major cell classes (glia, neurons) and separated out inhibitory and excitatory neurons using Seurat clustering and known marker gene enrichment.

### Defining cell types

Cell types were defined separately among inhibitory and excitatory neurons using the same procedure. First, SCTransform^93^ was used to normalize all C57BL6/J data across all ages, regressing out percentages of mitochondrial UMIs, ribosomal UMIs, and largest gene. Each cell from this dataset was then mapped to the published POA reference atlas^1^, separately for scRNA-seq and MERFISH reference atlases, using canonical correlation analysis-based label transfer^36^. Both reference datasets were normalized using SCTransform, and FindTransferAnchors was run using the SCTransform assays, PCA as the reference reduction, and 50 PCs. TransferData generated predictions to reference atlas cell types using 50 PCs. Each cell was then assigned (1) a scRNA-seq and (2) a MERFISH reference atlas predicted cell type, if the top prediction score exceeded 0.6; otherwise, a predicted cell type label was not assigned. These predicted cell type labels were used to guide subsequent clustering and cell type definition.

Then, P65 neurons were subset out in order to initially define cell types in adult neurons. Seurat functions RunPCA and FindNeighbors were run (150 PCs) and FindClusters (resolution 12) was used for an initial round of clustering. This generated clusters that often, but not always, had clear matches to POA reference atlas clusters, in terms of (1) majorities of cells predicted as a single MERFISH and/or scRNA-seq cell type (as described above) (Extended Data Fig 1b), with scRNA-seq and MERFISH cell types matching as described in the POA atlas paper^1^; and (2) marker genes matching those described in the POA atlas paper^1^. Each cluster was assigned a cell type ID, with occasional grouping of 2-4 clusters that mapped to identical sets of reference atlas cell types. Grouping of clusters into sub-regional groups was determined by correlation analysis (Fig 1h and Extended Data Fig 4d) and prior knowledge about spatial location from MERFISH (Moffitt et al., 2018^1^ fig 5c and S18). In cases where cell types lacked clear matches to MERFISH data, we estimated location based on three-color RNA scope staining and Allen Brain ISH Atlas data.

Having defined cell types at P65, we next mapped younger ages iteratively onto older ones (P28 to P65, then P18 to P28+P65, etc.). For this, we subset each younger age, processed those data as described above for P65, then used FindTransferAnchors and TransferData as described above but with 150 PCs and with older ages as the reference atlas, to generate prediction scores for each cell. Cells with top prediction score exceeding 0.8 were assigned to that cell type ID. Cells with top prediction score exceeding 0.5 were also assigned to that cell type ID, as long as the prediction score was at least twice as high as the 2^nd^-best prediction score. We chose these thresholds by exploring how well de novo clustering at each younger age generated clusters with homogenous predicted cell type ID labels. This procedure generated cell type IDs for 90%+ of cells at older ages, with lower percentages around P0 and earlier, which could be due to less precise dissection of smaller brains and thus inclusion of cells from brain areas not included at older ages. Prediction scores for cells that passed these thresholds are plotted in Fig 1e-f, averaged within a cell type. Identity ratio in Fig 1e is calculated using matrices as in Fig 1f: the value on the diagonal (best cell type match) minus the top off-diagonal value (second-best cell type match), divided by the value on the diagonal. Cell types in mutant and control samples were determined using the same procedure, except that cells were label transferred onto all ages. This procedure assigned cell type IDs to 80-90% of cells in these samples, an accuracy similar to across-age mapping in C57BL6/J samples. The remaining 10-20% of cells not assigned IDs in this manner were removed from further analysis.

To identify cell types in visual cortex at P10 from Trpc2 mutants and control littermates, we took a similar approach, using P8 and P14 visual cortex datasets as a reference atlas from ref. ^7^. We normalized P8 and P14 visual cortex data using SCTransform, used 150 PCs to perform label transfer, and used cutoffs as described above to assign predicted cell type IDs. 93% of cells in our samples were assigned predicted cell type IDs by this procedure.

### Regionalization quantifications

Matrices in Fig 1g-h and Extended Data Fig 4c-d were calculated by averaging scaled RNA values among all cells within a cell type, then calculating the Pearson correlation between all such cell type vectors. Correlations among cell types within a region were averaged to generate Fig 1i and Extended Data Fig 4e. Seurat’s FindAllMarkers function was used to identify regional marker genes in Extended Data Fig 4f, using an adjusted p-value cutoff of 0.05.

### devDEG analysis

The devDEG analysis presented in Fig 1j-m (and used to calculate heatmap data in Exended Data Fig 7e-f and Refinement Score in Fig 3e) comprises a combination of R and Python scripts from ref. ^40^ (https://zenodo.org/records/7113422) that we adapted. Each cell type was subset across all ages, then sample-pseudobulked (n=4 samples per age, except n=2 at E16) and passed to the limma-voom pipeline^94^ from the edgeR package^95^ for differential expression analysis testing. To implement this, a python 3.10.9 conda environment was created to install and isolate all necessary packages. We subset Seurat objects by cell types (inhibitory, excitatory) using Seurat, then converted them first to H5 format (with SaveH5Seurat) and then to h5ad format for import in scanpy (with the Convert function from the SeuratDisk package https://github.com/mojaveazure/seurat-disk). We created celltype – data point batches using 02.pseudo-bulk-by-batch_limma-voom_data-prep.ipynb notebook. We detected DEGs with edgeR/limma-voom using 10_major_trajectory_devDEG.R R-script, defined as those with FDR < 5%. We fit linearGAM trends (Generalized Additive Models) with LinearGAM (https://pygam.readthedocs.io, 11 dev-DEGs_age-trend-fits_rate-of-change.ipynb) for every gene across time points. We plotted trends using matplotlib (12 major-trajectory_dev-DEGs_stage-trend-fits.ipynb). We performed hierarchical clustering of DEG trends with 14 stage-trend-fits_clustering.ipynb and plotted the figures with matplotlib. This step was memory intensive and required up to 210G of RAM for the largest dataset in our analysis. For each DEG cluster we performed Gene Enrichment Analysis (15 Trajectory-devDEG_Gos-terms.ipynb). We plotted cell type / time point expression heatmaps for genes in GO categories (16 Eigentrends-of-selected-GO-terms.ipynb).

For Fig 2l, we used a combination of GO terms and the SynGO knowledge base on biological process ontology terms (ref ^96^, Supplementary Table S2, Biological process tab) to highlight expression values in GO categories in our dataset. We selected all categories that included the terms “synapse”, “synaptic”, “dendritic”, “neurotransmitter” and other key words to capture relevant categories in the SynGO knowledge base. A gene ontology term is represented by a set of genes. We overlapped DEG detected in each cell type across time points with genes in the selected GO category, and plotted expression heatmaps for categories where the median number of overlapping genes was >=40 when calculated across all cell types.

### Developmental trajectory quantification

To calculate developmental trajectories as in Fig 2a, we followed the approach described in ref. ^41^, to calculate distance between centroids in high-dimensional PC space. We subset each cell type across all ages, performed SCTransform to normalize and identify 2000 highly variable genes, and performed PCA (100 PCs). We then decided how many PCs to use for each cell type by asking how many of the top PCs are needed to explain 20% of variance in the data; if over 100 PCs were needed, we used 100 PCs. We then found the centroid at each age for each PC in gene expression space, and calculated the Manhattan distance between each centroid and the P65 centroid. This gives us a distance value for each age relative to P65. This distance generally decreased with age, indicating maturation, but on occasions where it increased, we set the distance value at the later age to that of the previous age. This step partly excludes transient movement away from adult (e.g. the transient up-regulation of gene expression programs related to axon guidance, which may be low at early and late ages but high at intermediate ages), but allowed us to better quantify how “adult-like” each cell type is at each age.

We additionally performed nearest neighbor analysis as an alternative measure of maturity state (Extended Data Fig 6e and 8a), following the method described in ref. ^40^ and associated code downloaded from https://zenodo.org/records/7113422. We focused on each cell type individually and constructed k-nearest neighbors (KNN) graphs that included all age categories, with each graph comprising 50 neighbors. To assess the maturity level for each age group, we calculated the fraction of nearest neighbors that originated from adult nuclei. This involved tallying the number of adult nuclei serving as nearest neighbors to the nuclei of a specific age. Subsequently, we normalized this total by the overall count of nearest neighbors, thereby deriving a proportion that reflects the maturity state based on the presence of adult nuclei among the nearest neighbors.

### Signaling gene expression analysis

All gene expression quantifications plotted were performed using log-normalized counts (i.e. Seurat’s “RNA” assay, data slot). We compiled lists of genes belonging to various signaling classes (Supplementary Table 2, based on refs. ^97,98^) and computed module scores using Seurat’s AddModuleScore function. Module scores were averaged among cells within an age and then z-scored across age. Refinement Score was calculated for each gene set by first summing the number of significant devDEGs (see “devDEG analysis” above) between P10 and P65 and then normalizing by dividing by the number of gene x cell-type combinations (after excluding combinations for which no expression is found at any age).

NeuronChat analysis^53^ was performed following tutorials from https://github.com/Wei-BioMath/NeuronChat. Ligand-receptor interactions present in the hypothalamus but not in the original interaction database were added utilizing The IUPHAR/BPS Guide to Pharmacology^98^ (Supplementary Table 3), and then all links that were not neuropeptide and monoamine links were removed. run_NeuronChat was run using M=100, mean_method=”mean”, and fdr=0.01. Links with ligand abundance less than 0.025 were filtered out.

### Distance analysis to determine male-female or mutant-control differences

We adapted a distance metric analysis from ref. ^41^ (also used in ref. ^99^) to ask whether male and female (or mutant and control) samples cluster in distinct regions of high-dimensional PCA space, separately for each cell type. To calculate distances between sample centroids, we used procedures and parameters similar to those described above for developmental trajectory calculation. We subset each cell type and, including both male/female or mutant/control samples, performed SCTransform to normalize and detect 2000 highly variable genes, and performed PCA (100 PCs). We then decided how many PCs to use for each cell type by asking how many of the top PCs are needed to explain 20% of variance in the data; if over 100 PCs were needed, we used 100 PCs. Then, for each sample (e.g. 4 male and 4 female samples), we quantified the sample’s centroid in PC space. We then calculated the Manhattan distance between all pairs of sample centroids. Our effect size (“Male-female PCA distance, normalized” or “Mutant-control PCA distance, normalized”) was calculated as the average across all male-to-female or mutant-to-control distances, normalized by the average of intra-sex or intra-genotype distances. This ratio quantifies how separated genotypes or sexes are compared to separation within each group. We calculated a two-sided t-test p-value between all inter-sex vs. intra-sex distances, or all inter-genotype vs. intra-genotype distances. In Fig 4f-g and Extended Fig 8b,c, any raw p-value >0.05 was considered Not Significant (N.S.), and results were validated by sexDEG analysis. For Fig 5a and Extended Data Fig 9a, Benjamini-Hochberg multiple comparisons corrected adjusted p-values were used to determine which cell types were Not Significant (N.S.). For sex analysis, to increase our n number, we included control samples from mutant/control experiments. For mutant analysis, to increase our n number, we included C57BL6/J samples as controls at the appropriate age, and confirmed minimal detectable differences between these samples and littermate control samples. We did not collect heterozygous control littermates for Trpm8 animals, and instead used same-age C57BL6/J samples as controls.

### sexDEG analysis

To determine DEGs between sexes, we performed sample-pseudobulk-based DESeq2 differential expression testing for each cell type. After removing mitochondrial, ribosomal, Y chromosome, and X inactivation genes, we split the dataset by ages. Then, we aggregated raw counts from cells of the same sample to use as input for DESeq2 and shrunk the log2 fold changes using lfcShrink (type=”apeglm”) to generate lists of differentially expressed genes between either male and female samples (sexDEGs). DEGs with adjusted p-value less than 0.05 were considered significant and used for downstream analysis. To increase our n number, we included control samples from mutant/control experiments. For GO term analysis we used gprofiler2 in R.

### Augur classifier analysis

To identify which celltypes have a notable change in expression between the control and mutant conditions, we used the package Augur v1.0.3, which performs cross-validated random-forest classifier analysis of single-cell datasets. First, we split the dataset by age and removed mitochondrial and ribosomal genes. Then we applied Augur (var_quantile=0.9, subsample_size=20) to generate AUC scores. For this analysis, control data included only control littermates with no added C57BL6/J samples (except for Trpm8 mutants, where C57BL6/J samples were the only available controls).

### Pseudotime analysis of Trpc2 mutant and control datasets

For each cell type, pseudotime was calculated across all ages from C57BL6/J data using Slingshot^73^ on log-normalized gene expression data and the top 20 PCs calculated from 2000 variable features. Then, for each mutant or control Trpc2 sample, cells from that sample were projected into the C57BL6/J 20-PC space (using stats::predict) and Slingshot prediction was performed (using slingshot::predict) to give each cell a predicted pseudotime value. Pseudotime values were averaged across all cells within each sample, and then between samples, to give data shown in Fig 5c. Two-sided t-test was performed to calculate p-value between control and mutant cell type pseudotime values.

#### Single-nucleus ATAC-sequencing analysis

snATAC-seq profiles were filtered to include only nuclei with at least 500 fragments, and only nuclei with paired snRNA-seq profiles passing QC as described above. C57BL6/J samples were split into two replicates, with each replicate containing 1 male and 1 female sample at each age. Peaks were called for each cell type in each replicate using macs2 callpeak command with parameters ‘‘–shift -100 –extsize 200 –nomodel –callsummits –nolambda – keep-dup all -q 0.05”. Only reproducible peaks that are present in both replicates were kept (n=778,573 peaks). Finally, to compile a union peak set, we combined peaks from all cell types and extended the peak summits by 200 bp on either side. Overlapping peaks were then handled using an iterative removal procedure. First, the most significant peak, i.e., the peak with the smallest p value, was kept and any peak that directly overlapped with it was removed. Then, this process was iterated to the next most significant peak and so on until all peaks were either kept or removed due to direct overlap with a more significant peak. Differentially accessible peaks were identified using the getMarkerFeatures() function from the ArchR package using a Wilcoxon rank sum test and accounting for bias introduced by TSSEnrichment and log10(nFrags). Peaks with FDR <= 0.1 and log2FoldChange >= 0.25 were considered cell-type-specific. Known transcription factor binding motifs present in the “vierstra” motifset were assigned to differentially accessible genomic regions and motif enrichment was evaluated using a hypergeometic test implemented by the peakAnnoEnrichment() function. Enriched motifs were those with minimum p-adjusted enrichment = 20, top 3 displayed per cell type (Extended Fig 5b). Motif footprints were calculated by combining all peaks harboring a given motif in aggregate and accounting for Tn5 insertion bias via the getFootprints() function.

#### Statistics and reproducibility

Data were processed and analyzed using a combination of R and Python codes. Sample sizes were chosen based on common practice in single-nucleus sequencing experiments. Individual data points were plotted wherever possible. Boxplots represent the median, first and third quartiles (hinges), and 1.5 * inter-quartile range (whiskers). Outliers are shown wherever individual data points are not plotted. All data was analyzed using two-tailed non-parametric tests. Statistically not significant is indicated by N.S., and significance was indicated by *P < 0.05, **P < 0.001, and ***P < 0.0001. Statistical details are given in the respective figure legends.

## Data Availability

The data generated in this study are available at https://data.nemoarchive.org/biccn/grant/u19_huang/dulac/transcriptome/sncell/10x_multiom e/

## Code Availability

Custom code used in this study is available upon request.

## Acknowledgements

We thank members of the Dulac laboratory, the Bauer Core Facility, Nathan Zemke, Jia Yin Xiao, Dhananjay Bambah-Mukku, Will Allen, and Jenny Chen for helpful advice on experiments, analysis, and the manuscript; Patrick Arter for assistance with RNA scope experiments; Salwan Butrus for providing data on visual cortex; and Renate Hellmiss for help with figures. This work was supported by Jane Coffin Childs Medical Research Award 61-1749 to H.K., NIH award K99HD108801 to B.L.L., NIH grant 5UM1HG011585 to B.R. and C.D., NIH grants U19MH114821, R01HD082131, and R01NS112399, and a NOMIS Foundation Award to C.D. C.D. and D.D.G. are investigators at the Howard Hughes Medical Institute.

## Author Contributions

H.S.K. and C.D. conceived and designed the study. H.S.K. performed multiome sequencing experiments and analyzed snRNA-seq data with help from N.S. and S.N. for DEG and classifier analysis and K.Z. for nearest neighbor maturation analysis. B.L. and K.Z. analyzed snATAC-seq data. C.S. raised and provided mice for touch mutant experiments. C.D., B.R., D.D.G., and S.J.H.S. provided instruction and comments during the research. H.S.K. and C.D. wrote the manuscript with input from all authors.

